# Selective Elimination of TP53 Mutant Cells by Transcript-Activated Chromatin Shredding

**DOI:** 10.64898/2026.05.08.723607

**Authors:** Jingkun Zeng, Zhiyuan Cheng, Huadong Chen, Jared Thompson, Kadin T. Crosby, Hesong Han, Wayne Ngo, Chenglong Xia, Daniel Rosas-Rivera, Min Hyung Kang, Ying Mao, Giselle Lee, John F. X. Diffley, Yixuan Song, Longhui Qiu, Nathan M. Krah, Niren Murthy, Ryan N. Jackson, Yang Liu, Alan Ashworth, Jennifer A. Doudna

## Abstract

Genetic mutations that drive cancer often occur in tumor suppressor proteins, including the p53 transcription factor which is altered in ∼40-50% of cases^1,2^. However, current therapies fail to target most such mutations because the mutant proteins typically lack defined drug-binding pockets, and restoring the endogenous function has proven challenging. Here, we programmed CRISPR-Cas12a2, an RNA-guided nuclease with *trans*-nucleolytic cleavage activities^3,4^, to selectively kill cancer cells by targeting cancer-specific transcripts. This approach eliminates cells by inducing *trans* chromatin cleavage, triggering DNA damage and cell death. Unlike existing methods, RNA-guided Cas12a2 senses cellular RNA signatures to shred chromatin, enabling precise targeting of undruggable mutations. Transcript-activated chromatin shredding provides an innovative paradigm to develop precision disease treatments for undruggable targets.

---

CRISPR-Cas12a2 is an RNA-guided nuclease that cleaves both RNA and DNA in *trans* upon target RNA recognition^3,4^. In bacteria, phage or other foreign transcripts trigger Cas12a2-guide RNA (gRNA) catalyzed depletion of cellular nucleic acids to induce cell dormancy or death. This abortive infection mechanism is thought to prevent phage propagation within the bacterial community by killing just those cells expressing a Cas12a2-guide RNA-detectable transcript.

The field of cancer biology has long appreciated the utility of targeted cell killing because tumor cells express aberrant or mutated proteins not found in healthy tissue. *TP53,* encoding the most common cancer-associated transcription factor p53, is mutated in ∼40-50% of all cancers and up to 70–90% of ovarian, NSCLC (non-small cell lung cancer) and pancreatic tumors^1,2,5^. *TP53* mutations also tend to be clonal, arising early and persisting within a heterogenous population of tumor cells^6,7^. Restoring p53 function for tumor regression has been considered the “holy grail” of cancer therapy^8–13^. However, no approved therapies are available to target the p53 protein due to its lack of druggable pockets and the difficulty of activating defective transcription factors. Furthermore, a potential side effect of p53 activators is that unintended p53 activation in healthy cells can induce senescence and whole-genome duplication^14–16^, complicating therapeutic strategies. An alternative would be just killing the cells with mutant proteins.

Here, we explored the potential for CRISPR-Cas12a2 to selectively eliminate mammalian cells expressing specific mRNAs, including mutant *TP53* transcripts. We found that, upon RNA-guided RNA target recognition, Cas12a2 cleaves eukaryotic chromatin in *trans*, triggering DNA damage responses and cell death in mammalian cells. We show that the mRNAs encoding mutant proteins can be targets for triggering for Cas12a2-mediated killing. In particular, guide RNAs targeting single nucleotide variants (SNVs) were designed to selectively eliminate cells expressing several *TP53* point mutations. We show that this approach is highly specific, causing DNA damage-induced cell death only in the presence of the mutant but not the wild-type transcript. Delivering Cas12a2 mRNA and its guide RNA targeting c-MYC and p53 R248Q transcripts using lipid nanoparticles (LNPs) reduced the tumor burden *in vivo*. Together, these data suggest that cancer-specific cell targeting by transcript-activated chromatin shredding could be a valuable approach to cancer therapy.

## Results

### Cas12a2 shreds chromatin in trans upon RNA targeting in vitro

We first examined Cas12a2’s *trans* cleavage activity *in vitro* using purified SuCas12a2 ribonucleoproteins (RNPs) with fluorescently labelled linear nucleic acid substrates. SuCas12a2 RNPs specific for the target RNA degraded both FAM-labelled RNA and dsDNA substrates (Fig. 1a). SuCas12a2 RNPs also cleaved supercoiled plasmid DNA *in trans* following RNA targeting, whereas control SuCas12a2 RNPs containing non-targeting guide RNAs did not induce RNA or DNA cleavage (Fig. 1b). These results are consistent with previous work showing that Cas12a2 degrades naked nucleic acids *in trans*^3,4^.

**Fig 1.**
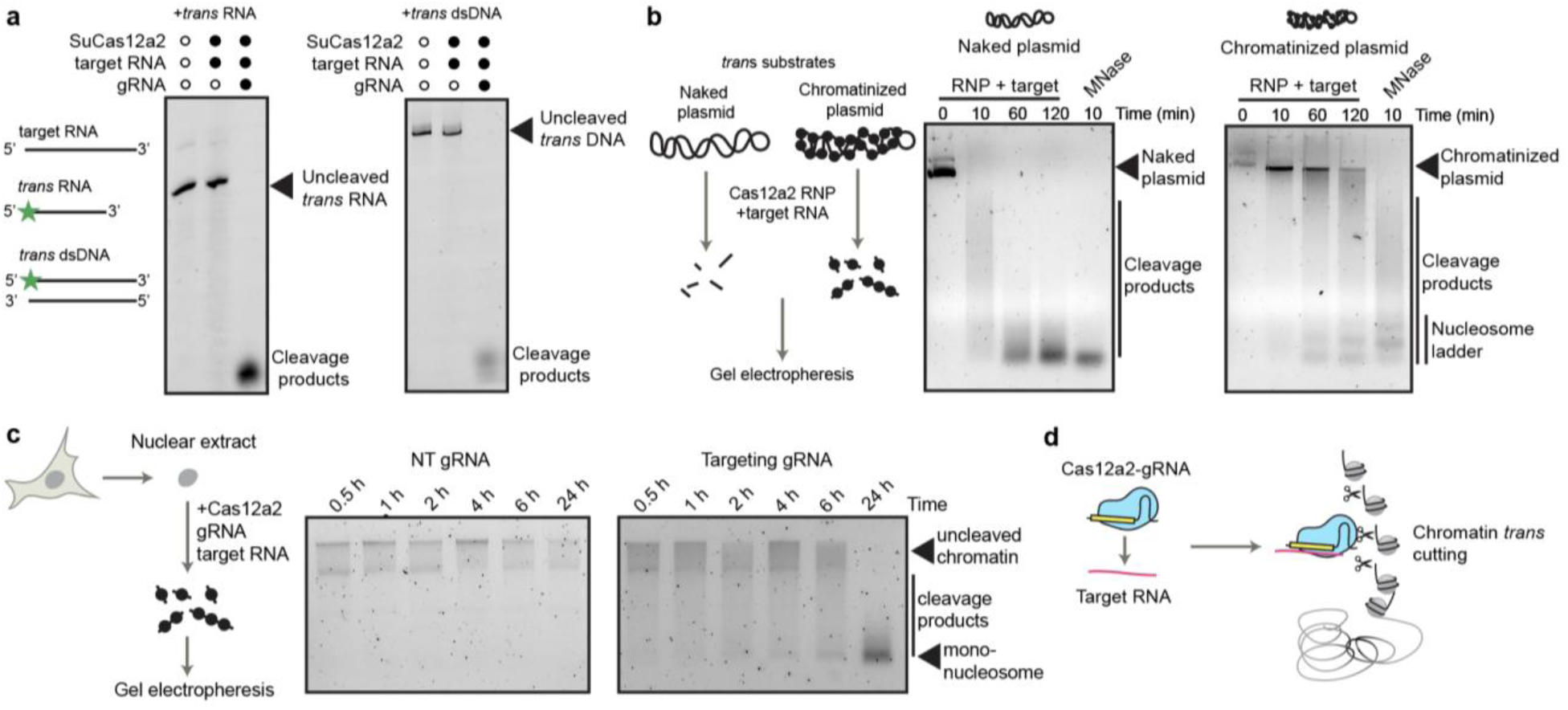
Biochemical analysis of chromatin *trans*-cleavage by Cas12a2. **a,** *Trans*-cleavage of FAM-labelled non-target ssRNA and dsDNA substrates by purified Cas12a2 protein. **b,** Time-course analysis of *trans*-cleavage of a naked supercoiled plasmid and a chromatinized plasmid by Cas12a2-gRNA complex incubated with a target RNA. **c,** Time-course analysis of *trans*-cleavage of chromatin in HEK293 nuclear extracts by Cas12a2 incubated with a target RNA as well as a non-targeting (NT) gRNA or a targeting gRNA. **d,** Schematic showing chromatin *trans*-cleavage by Cas12a2 upon RNA targeting.

We next tested whether Cas12a2 RNPs can cleave chromatinized DNA, as would occur in eukaryotic cell nuclei. A 10.6 kb chromatinized plasmid was gradually degraded *in trans* by SuCas12a2 RNPs upon RNA targeting, although at a slower rate than was observed for the naked plasmid (Fig. 1b). A distinct ladder of DNA cleavage products formed after 1 h of incubation with SuCas12a2 RNPs, corresponding to the sizes of mono-, di- and tri-nucleosomes, suggesting SuCas12a2 preferentially cleaved linker regions between nucleosomes. To test whether Cas12a2 RNPs can also degrade mammalian chromatin, we incubated nuclei extracted from HEK293 cells with SuCas12a2 RNPs, and a target RNA. Whereas RNPs containing a non-targeting (NT) gRNA induced no observable cleavage, RNPs containing a gRNA with sequence complementarity to target RNA led to degradation of HEK293 chromatin (Fig. 1c). These results suggest that Cas12a2 can serve as an RNA-guided chromatin shredder (Fig. 1d).

### Transcript targeting by Cas12a2 RNPs kills mammalian cells

RNA-guided chromatin shredding could potentially induce programmed killing of mammalian cells. To test Cas12a2 activation in mammalian cells, we nucleofected different doses of SuCas12a2 RNPs into HEK293T cells stably expressing green fluorescent protein (GFP) (hereafter HEK293-GFP) (Fig. 2a), targeting GFP mRNA transcripts. Live cell imaging showed that almost no cell proliferation occurred in HEK293-GFP cells nucleofected with 10-100 pmol SuCas12a2 RNP containing GFP-complementary gRNA (GFP gRNA RNP) (Fig. 2a-c; Extended Data Fig. 1a). The treated cells contained enlarged or fragmented nuclei after 72 h, indicative of mitotic bypass or mitotic catastrophe after extensive DNA damage^16^ that could lead to a loss of cell proliferation capacity (Fig. 2b). These cells also showed increased expression of γH2AX, a marker for DNA double-stranded breaks, and phospho-KAP1 (S824), a marker for heterochromatic DNA damage^17^ (Fig. 2d), consistent with Cas12a2-mediated chromatin *trans*-cleavage.

**Fig 2.**
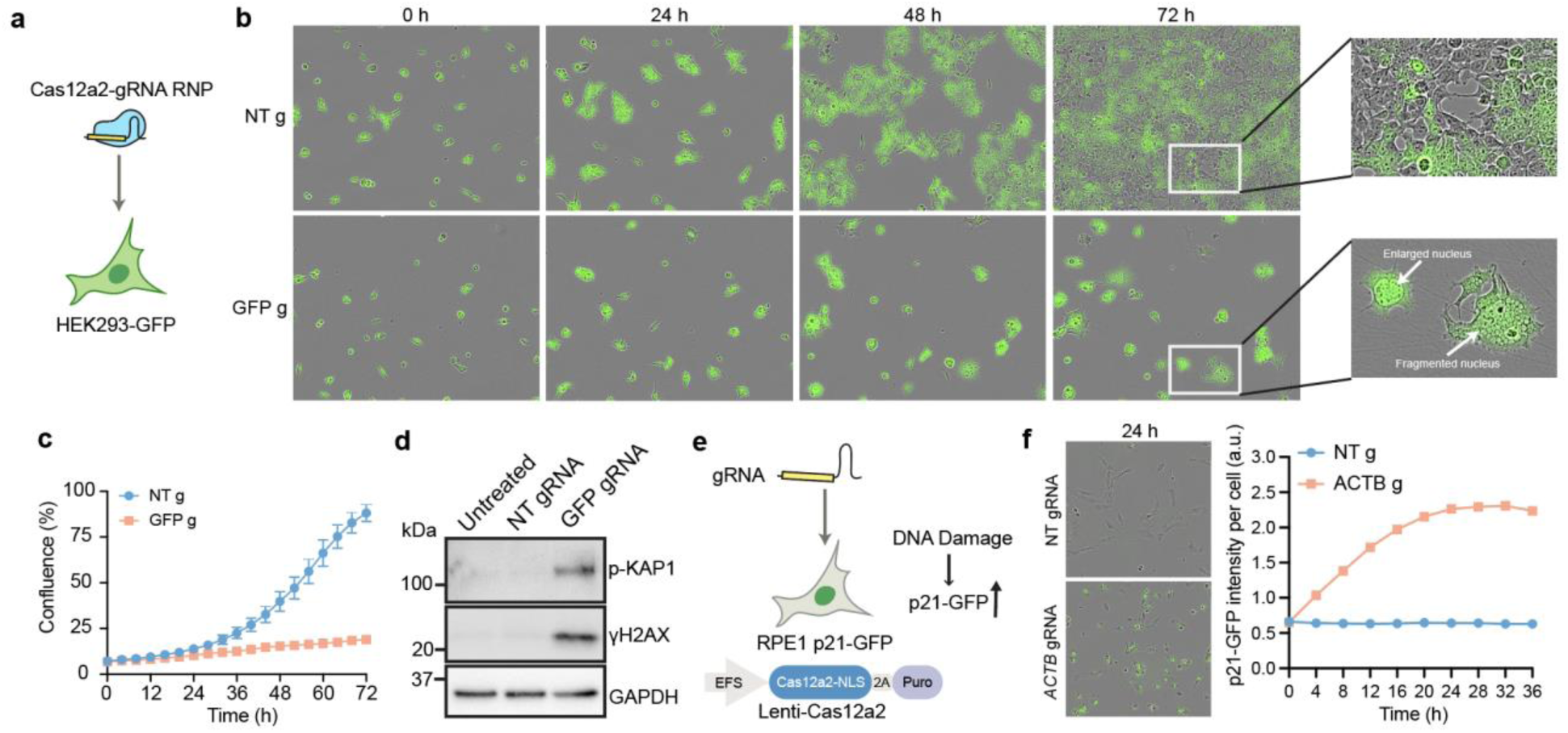
Cas12a2 induces acute DNA damage and cell cycle arrest upon RNA targeting in mammalian cells. **a,** Nucleofection of Cas12a2-gRNA ribonucleoprotein (RNP) complexes into HEK293 cells expressing GFP. **b,** Representative time-lapse images of HEK293-GFP cells nucleofected with Cas12a2-gRNA RNPs with a NT gRNA (NT g) or GFP-targeting gRNA (GFP g). Merged green and phase contrast channels are shown. **c**, Cell confluence measurement of images taken every 4 h in **b. d,** Immunoblots showing expression of DNA damage markers in HEK293-GFP cells 48 h following Cas12a2-gRNA RNP nucleofection. **e,** A genotoxic stress reporter RPE1 cell line expressing GFP-tagged DNA damage response factor p21, and transduced with Cas12a2-NLS under an EFS promoter. 2A: self-cleaving peptide. Puro: puromycin resistance gene. **f,** Time-lapse images following transfection of an *ACTB*-targeting gRNA. Representative images at 24 h and increases in GFP signals over time are shown.

To investigate whether target transcript abundance affects Cas12a2-induced cell death, we established single cell clones of HEK293T cells expressing high, medium and low levels of GFP (referred to as HEKGFP High, HEKGFP Mid and HEKGFP Low) (Extended Data Fig. 1b). Using single-molecule fluorescence *in situ* hybridization (smFISH), we quantified GFP mRNA transcripts in these cells, detecting ∼550 transcripts per cell in HEKGFP Low, ∼1,100 in HEKGFP Mid, and an unquantifiable number in HEKGFP High due to image saturation (Extended Data Fig. 1c, d). Following nucleofection with a low dose (5 pmol) GFP gRNA SuCas12a2 RNP, HEKGFP High cells showed almost no cell proliferation, whereas HEKGFP Mid and HEKGFP Low showed milder growth defects (Extended Data Fig. 1e). Control cells with no GFP expression were unaffected. These results suggest that the extent of Cas12a2-mediated cell death in mammalian cells correlates with target transcript abundance.

### Cas12a2 activity depends on Mg^2+^ concentration

Previous biochemical assays using Cas12a2 RNPs were performed in the presence of 10 mM MgCl_2_ ^3,4^, whereas free magnesium ion concentration in mammalian cells is ∼0.1-1 mM^18^. CRISPR-Cas9 enzyme activity is known to be slower at mammalian cellular Mg^2+^ levels^19^, so we asked whether Cas12a2 is similarly affected by Mg^2+^ concentration. To assess the efficiency of Cas12a2 cleavage under conditions similar to those of mammalian cells, we performed *in vitro trans* cleavage assays over a range of Mg^2+^ concentrations. Although Cas12a2 RNP activity decreased at lower Mg^2+^ concentrations, DNA cleavage in *trans* occurred even at 0.1 mM Mg^2+^ (Extended Data Fig. 1f).

We next tested whether Mg^2+^ concentration affects RNP formation or cleavage activity. We pre-incubated SuCas12a2 protein with GFP gRNA at Mg^2+^ concentrations ranging from 0.1 mM to 10 mM, followed by nucleofection into HEK293-GFP cells. Regardless of the Mg^2+^ concentration used for RNP pre-incubation, cells were induced to stop growing with the same efficiency (Extended Data Fig. 1g). These results suggest that despite being slower under mammalian cell conditions compared to conditions in bacterial cells, Cas12a2 RNP activity is sufficient to induce mammalian cell death.

### Cas12a2 activation triggers an acute DNA damage response in mammalian cells

HEK293T cells express SV40 large T antigen which inhibits key cell cycle and DNA repair regulators RB (retinoblastoma) and p53^20,21^. To examine Cas12a2-induced stress responses in a more normal mammalian context, we used hTERT-RPE1 (hereafter RPE1), an untransformed epithelial cell line immortalized by telomerase and bearing wild-type p53 activity. We stably expressed nucleoplasmin nuclear localization signal (NLS)-tagged SuCas12a2 in RPE1 cells under a EF1-a short (EFS) promoter, and knocked in GFP at the C-terminus of the p53 downstream cell cycle inhibitor *CDKN1A* (encoding p21) (Fig. 2e). This created a genotoxic stress reporter cell line, wherein GFP expression increases in response to DNA damage as a result of p21 expression.

Using this reporter cell line, we assessed stress responses upon Cas12a2 activation. Transfecting gRNAs targeting *ACTB* (encoding β-ACTIN) or *GAPDH* transcripts led to cell growth arrest and expression of DNA damage markers (Extended Data Fig. 2a, b). Among the seven *ACTB* gRNAs tested, the ones that produced the strongest cell killing also caused the largest increase in GFP signal (Extended Data Fig. 2c), consistent with higher levels of DNA damage. GFP expression increased within 4 hours after transfecting *ACTB* gRNAs, peaking at around 24-48 h (Fig. 2f), suggesting Cas12a2 cleaved chromatin shortly after target transcript recognition in mammalian cells to trigger DNA damage responses.

### Transcript thresholds enable selective Cas12a2 killing of oncogene expressing cells

Elevated expression of oncogenes is a common driver of tumorigenesis. For example, *CCNE1* and *MYC* are frequently amplified in various cancers^22^, leading to high transcript levels (Extended Data Fig. 3a, b). Since higher target expression levels correlate with more efficient Cas12a2 RNP-induced cell killing (Extended Data Fig. 1b-e), we reasoned that Cas12a2 RNPs might selectively kill cancer cells expressing high levels of oncogenes. To explore this, we used U2OS cells expressing doxycycline (Dox)-inducible cyclin E1 (hereafter U2OS TetOn CycE) (Extended Data Fig. 3c), which over-express cyclin E1 at a level comparable to patient-derived cancer cells^16,23^. Before and after Dox treatment, there were ∼50 and ∼560 *CCNE1* transcripts per cell respectively (Extended Data Fig. 3d). We tested six gRNAs targeting the *CCNE1* mRNA transcript in U2OS TetOn CycE cells stably expressing Cas12a2. Whereas some gRNAs (gRNA 3, 4 and 5) induced cell growth defects in both untreated cells and Dox-treated cells, gRNA 1 and 2 induced growth defects selectively in Dox-treated cells (Extended Data Fig. 3e). These observations suggested that a targeting window exists to distinguish between high and low levels of the same transcript. Taken together, these results imply that transcripts expressed at elevated levels in cancer cells can be targeted for Cas12a2-mediated cell killing.

### Targeting oncogene transcripts containing an in-frame deletion kills cells

In-frame insertion and deletion (indel) mutations can lead to hyperactivation of oncogenes. *EGFR* (epidermal growth factor receptor) exon 19 deletion mutations are common activating mutations found in NSCLC, making up ∼45% of all *EGFR* mutations^24,25^. The indel junction of such oncogene transcripts creates a sequence that does not exist in normal cells, which we reasoned can be targeted by Cas12a2 RNPs for selective cell killing. To test this, we used a human NSCLC cell line PC9 which contains an endogenous *EGFR* E746_A750 deletion (E746_A750del) mutation, resulting from a 15bp deletion in the *EGFR* gene (Fig. 3a, b). We designed a Cas12a2 gRNA that is complementary to the E746_A750del transcript (EGFRdel gRNA) around the deletion junction and should not anneal to the WT transcript. We tested whether it can selectively kill cells containing an *EGFR* E746_A750del mutation by using PC9 cells (harboring endogenous *EGFR* E746_A750del) and RPE1 cells (*EGFR* WT) stably expressing Cas12a2. Transfection of Cas12a2-expressing PC9 cells with the EGFRdel gRNA caused robust growth arrest and expression of DNA damage markers, but no effect was observed after similar transfection of RPE1 cells (Fig. 3b-e). This finding suggested that indel junctions in oncogenic transcripts can be targeted by Cas12a2 for selective cell killing.

**Fig 3.**
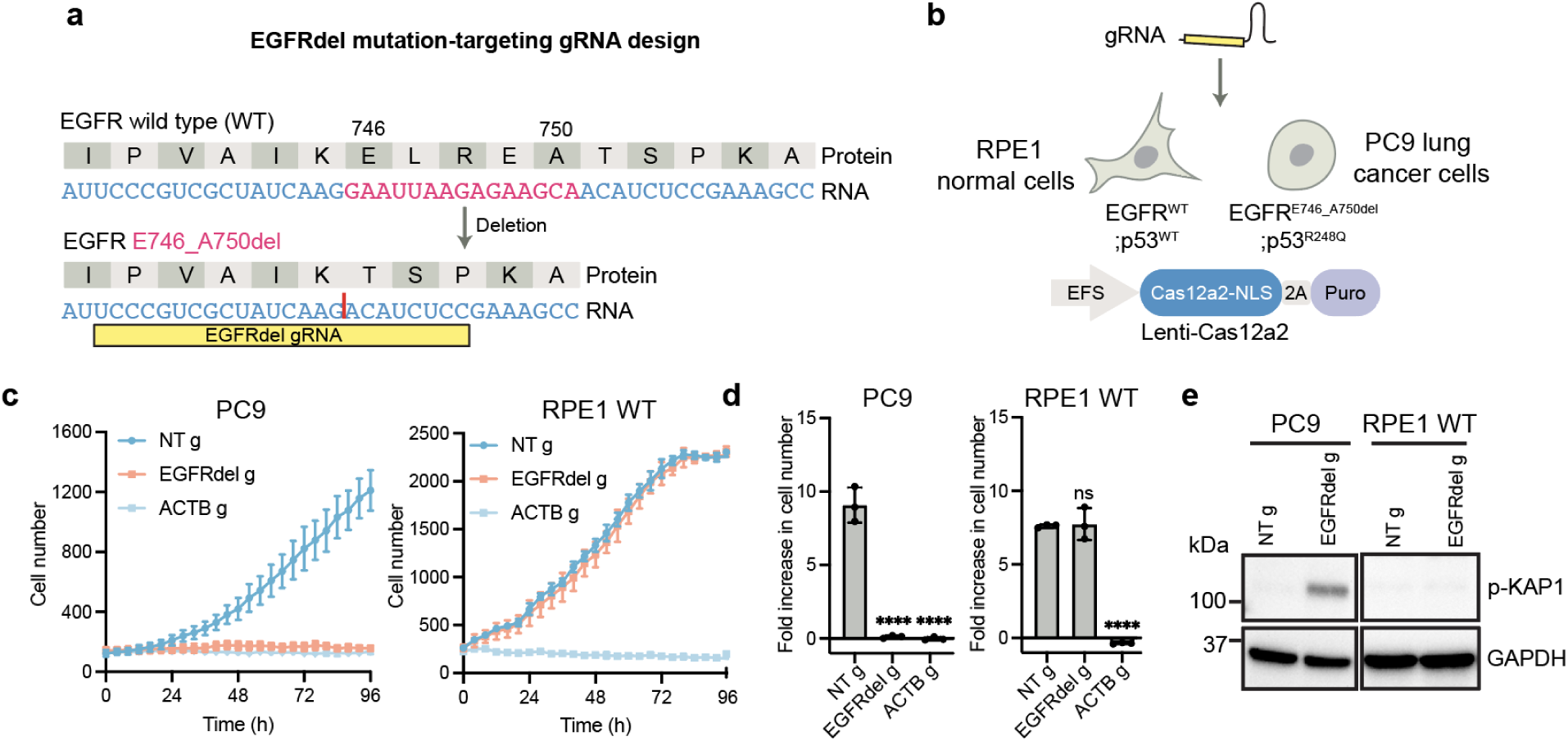
Selective killing of NSCLC cells harboring an *EGFR* in-frame deletion mutation. **a,** Design of a gRNA (EGFRdel g) targeting the EGFR E746_A750del mutant transcript. **b,** Testing of the EGFRdel-specific gRNA in PC9 NSCLC cells (EGFR E746_A750del) and RPE1 cells (EGFR WT) stably expressing Cas12a2. **c**, Growth curves of PC9 and RPE1 cells after targeting EGFR E746_A750del mutant transcript. **d**, Fold-increase in cell number over 96 h following Cas12a2 targeting of the EGFR E746_A750del mutant transcript in PC9 and RPE1 cells. **e,** Immunoblots showing expression of DNA damage markers in PC9 and RPE1 cells 48 h following Cas12a2 targeting of the EGFR E746_A750del mutant transcript. Statistics: ****p<0.0001; ns, non-significant, one-way ANOVA with Dunnett’s test.

### Targeting TP53 point mutations kills cells

Mutations in the p53 tumor suppressor protein, the most common driver for tumorigenesis (Extended Data Fig. 4a), often result from SNVs in the *TP53* gene sequence. Mutations in almost every residue of the p53 protein can contribute to carcinogenesis^26^, but certain ‘hotspot’ mutations are particularly common^27^ (Fig. 4a). For example, R248Q, caused by a G-to-A substitution in the gene sequence, makes up ∼7% of all *TP53* mutations. We wondered whether Cas12a2 can distinguish such SNVs in *TP53* mutant transcripts. SuCas12a2 recognizes an adenine-rich protospacer flanking site (PFS) adjacent to a targeted RNA sequence^3^ (Extended Data Fig. 4b), which functions analogously to the protospacer adjacent motif (PAM) in DNA-targeting CRISPR systems. We reasoned that the G-to-A mutation site in the R248Q mutant transcript could serve as a PFS (referred to as mutA PFS) to activate SuCas12a2 selectively (Fig. 4b). To test this, we stably expressed Cas12a2 and the p53 R248Q mutant transgene in RPE1 cells (RPE1 p53 R248Q). A gRNA was designed to anneal to p53 R248Q mutant mRNA next to the mutA PFS (R248Q gRNA). Transfecting this R248Q gRNA induced a robust growth arrest in RPE1 p53 R248Q cells, but no growth rate change in RPE1 WT cells (Fig. 4c and Extended Data Fig. 4c), indicative of selective Cas12a2 activation. To assess competitive growth between mutant and WT cells after Cas12a2 targeting, we co-cultured RPE1 p53 R248Q (expressing mCherry) and RPE1 WT cells (expressing GFP) at a 1:1 ratio at time zero (Fig. 4e and Extended Data Fig. 4d). Analysis 96 h after cell transfection with NT gRNA showed that R248Q cells outgrew WT cells, yielding a ratio of GFP/mCherry cell numbers of ∼0.62. In contrast, transfection with R248Q gRNA reversed this trend, increasing the ratio to ∼4.5 consistent with significant outgrowth of WT over R248Q mutant cells after targeting the mutant transcript.

**Fig 4.**
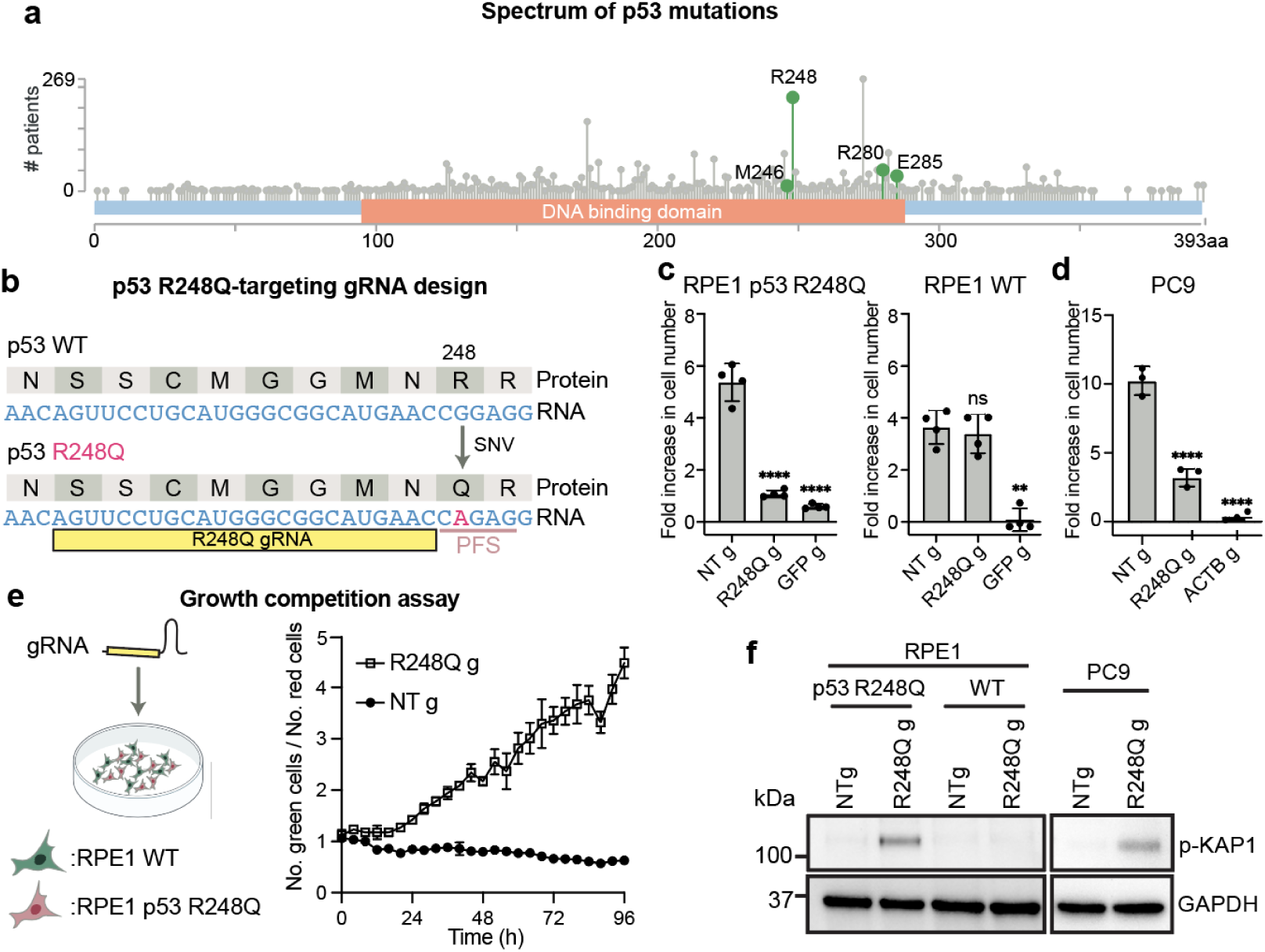
Selective killing of cells harboring *TP53* SNVs. **a,** Mutational spectrum of the p53 protein in tumor samples from 3,949 cancer patients from TCGA Pan-Cancer Atlas studies in the cBioPortal database. M246, R248, R280 and E285 residues targeted in this study are highlighted. **b,** Design of a gRNA (R248Q g) targeting the p53 R248Q mutant transcript. PFS, protospacer flanking site. **c,** Fold-increase in cell number over 96 h following Cas12a2 targeting of the p53 R248Q mutant transcript in RPE1 cells expressing p53 R248Q (RPE1 p53 R248Q), wild-type RPE1 cells (RPE1 WT) and **d,** PC9 cells. Cas12a2 was stably expressed in these cells. **e**, Growth competition assay between RPE1 WT and R248Q cells. **f,** Immunoblots showing expression of DNA damage markers in cells 48 h following Cas12a2 targeting of the p53 R248Q mutant transcript. Statistics: *p<0.05; **p<0.01; ****p<0.0001; ns, non-significant, one-way ANOVA with Dunnett’s test.

The results above used cells expressing p53 R248Q under a strong CMV promoter. To exclude transcript-level differences as a confounding factor, we tested R248Q gRNA in RPE1 cells expressing a control p53 mutation R175H also under the CMV promoter (RPE1 p53 R175H). At the R248 position, the p53 R175H transcript has the same sequence as p53 WT. We showed that transfecting R248Q gRNA did not cause a growth rate change in Cas12a2-expressing RPE1 p53 R175H cells (Extended Data Fig. 4c, e). Moreover, *TP53* transcript levels were similar in RPE1 p53 R248Q cells (∼400 per cell) and control RPE1 p53 R175H cells (∼500 per cell), suggesting the selectivity is not due to differences in expression levels, although RPE1 WT cells had fewer transcripts (∼25 per cell) (Extended Data Fig. 4f). We also detected increases in DNA damage markers (phospho-KAP1) only in RPE1 p53 R248Q cells (Fig. 4f and Extended Data Fig. 4g). Additionally, in biochemical assays, *trans* DNA cleavage by Cas12a2 RNPs containing R248Q gRNA was specific to the R248Q mutant transcript (Extended Data Fig. 5a). FUCCI (fluorescent ubiquitination-based cell-cycle indicator) live cell imaging^28^ showed that R248Q gRNA caused extended G2 arrest in RPE1 p53 R248Q cells, and an increase in the number of cells with fragmented nuclei, consistent with mitotic catastrophe after DNA damage (Extended Data Fig. 5b, c). These results indicate that Cas12a2 RNPs can precisely target *TP53* mutant cells with high selectivity and with minimal side effects in healthy cells.

To test whether cancer cells containing an endogenous p53 R248Q mutation can be targeted, we used PC9 cells that contain an endogenous p53 R248Q mutation (∼83 transcripts/cell) in addition to *EGFR* E746_A750del (Fig. 3a, b; Extended Data Fig. 4f). We observed that in Cas12a2-expressing PC9 cells, transfecting R248Q gRNA also caused significant growth defects and expression of DNA damage markers (Fig. 4d, f). The remaining cells tested positive with cell death stains (propidium iodide+ or Annexin V+) after 96 hours (Extended Data Fig. 5d, e). These results suggest that Cas12a2 RNPs can efficiently target cancer cells harboring endogenous *TP53* mutations.

Using similar design principles for R248Q gRNA, we designed gRNAs that selectively induced DNA damage and cell death in cells expressing p53 R280K and p53 E285K, both of which are caused by a G-to-A SNV in the *TP53* gene (Extended Data Fig. 6a-d). In addition, we screened six gRNAs to target the endogenous G-to-C SNV for the p53 M246I mutation in NCI-H23 cells (Extended Data Fig. 6e). Cas12a2 showed *trans*-cleavage selectivity *in vitro* with two of the gRNAs (gRNAs 4 and 6) targeting the M246I mutant transcript, suggesting a single mismatch between the gRNA-target duplex can decrease Cas12a2 activity (Extended Data Fig. 6f). Cas12a2 targeting with one of the selective gRNAs, M246I gRNA 6, induced growth defects in NCI-H23 cells (p53 M246I) but not in U2OS cells (p53 WT) (Extended Data Fig. 6g). Taken together, these results demonstrate that SNVs spanning *TP53* mutant transcripts can be exploited for Cas12a2-mediated selective cell killing.

To assess the fraction of *TP53* mutations targetable using Cas12a2, we analyzed *TP53* coding sequence mutations in 16,708 patient samples to identify potential Cas12a2 target sites. We found 25.7% of *TP53* mutations in patients are B-to-A substitutions (where B is G, C, or T) (Extended Data Fig. 6h), which could serve as new or enhanced PFS for targeting, as demonstrated with the R248Q mutation (Fig. 4b-d). Additionally, for 73.2% of *TP53* mutations in patients, there were A, ABA or AA motifs within 24 nt at the 3’ end (Extended Data Fig. 6i), which could serve as a PFS to allow Cas12a2 targeting.

### Co-delivering Cas12a2 mRNA and gRNA suppresses tumor progression in vivo

Towards the goal of eventually using Cas12a2 RNPs in therapeutic applications, we tested whether Cas12a2 can be delivered as mRNA for cell killing. We generated capped and pseudo-uridylated mRNA encoding NLS-tagged Cas12a2 by *in vitro* transcription (IVT). Co-transfection of PC9 cells with this Cas12a2 mRNA along with p53 R248Q gRNA induced significant growth defects compared to controls transfected with non-targeting gRNA (Extended Data Fig. 6j). Additionally, we co-packaged Cas12a2 mRNA and p53 R248Q gRNA lipid nanoparticles (LNPs)^29^ (Fig. 5a; Extended Data Fig. 6h). Addition of these LNPs to cultured PC9 lung cancer cells similarly induced robust growth defects compared to cells treated with LNPs prepared with non-targeting gRNA (Fig. 5b).

**Fig 5.**
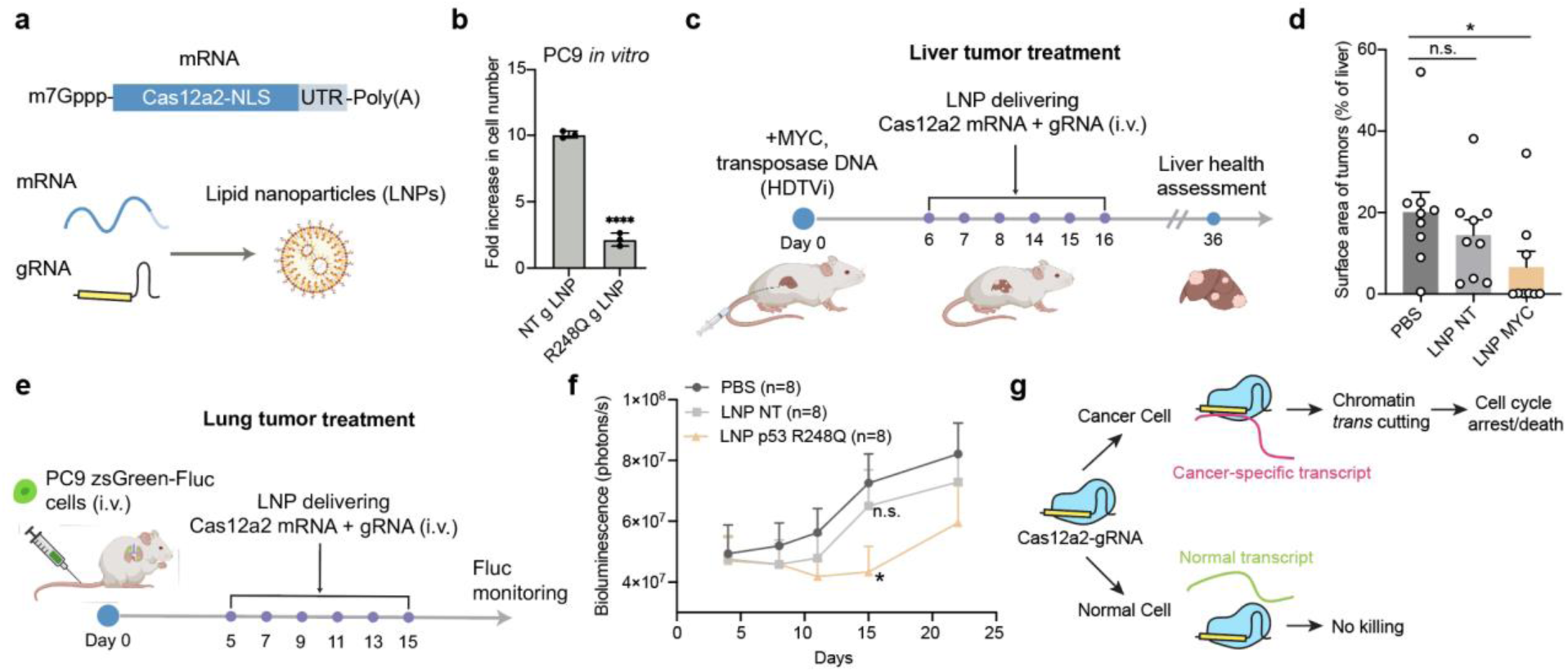
*In vivo* anti-tumor test. **a-b,** Cas12a2 targeting of p53 R248Q transcripts by delivering LNP packaging mRNA encoding Cas12a2 and gRNAs into PC9 cells. Fold-increase in cell number over 96 h is shown on the right. **c,** Schematic showing *in vivo* treatment of MYC-induced liver tumors. Plasmids expressing the MYC oncogene and Sleeping Beauty transposases were introduced into mice by hydrodynamic tail vein injection (HDTVi), resulting in stable integration of MYC into random liver cells. **d,** Quantification of liver tumor surface area as the percentage of whole liver surface area. **e,** Schematic showing *in vivo* treatment of PC9 lung tumors. The inoculated PC9 cells express firefly luciferase (Fluc) for live animal imaging. **f**, Quantification of tumor bioluminescence signals over time in treated mice. **g,** Schematic showing Cas12a2-mediated selective cell elimination. Statistics: *p<0.05; ns, non-significant, Student’s t-test.

To evaluate the therapeutic efficacy of Cas12a2 *in vivo*, we first used a previously described MYC-induced liver tumor model^30^. In this model, the MYC oncogene was stably integrated into random liver cells of tumor-prone FVB/NJ mice by transposases, resulting in liver tumor formation (Fig. 5c). We first screened gRNAs targeting MYC transcripts in Cas12a2-expressing HEK293 cells and identified one that induced strong DNA damage marker expression upon co-transfection with the MYC-expressing plasmid (Extended Data Fig. 7a). This gRNA (MYC g4) was subsequently co-packaged with Cas12a2 mRNA into LNPs for *in vivo* anti-tumor test (Fig. 5c). Treatment began on day 6 post-tumor induction. Mice receiving MYC-targeting LNPs exhibited reduced tumor surface area (percentage of total liver) and decreased liver-to-body weight ratios compared to control groups (Fig. 5d; Extended Data Fig. 7b-d).

Next, we evaluated p53 mutation targeting in mice bearing lung tumors. Lung-enriching LNPs have been previously developed^29,31,32^ via the SORT mechanism, with demonstrated delivery efficiencies of 15-20% for Cas9 editors in healthy lung tissues. In addition, SORT LNPs have also been able to deliver mRNA to lung metastases generated from A549 cells in mice^33^. We first assessed whether these LNPs could be used to deliver cargo to lung tumors generated from PC9 cells stably expressing zsGreen and a Cre-dependent tdTomato reporter (Extended Data Fig. 8a, b). Intravenous injection of PC9 cells established lung xenografts (Extended Data Fig. 8c, d). Following intravenous injection of a single dose of Cre mRNA-containing LNPs, ∼7-18% of zsGreen-labeled PC9 cells turned on tdTomato expression within the lung xenograft (Extended Data Fig. 8e-g). Although this delivery efficiency was modest, we proceeded with anti-tumor testing using multiple dosing to compensate for the limited cellular uptake.

We engrafted 200,000 firefly luciferase (Fluc)-expressing PC9 cells into immunodeficient mice and initiated treatment 5 days later with six doses of LNPs co-delivering Cas12a2 mRNA and gRNA (Fig. 5e). Tumor burden was monitored by bioluminescence imaging. Mice treated with p53 R248Q-targeting LNPs showed reduced tumor progression compared to control-treated groups (Fig. 5f). To assess therapeutic potential in a more challenging context, we established an advanced-stage tumor model by engrafting mice with 1 million PC9 cells and allowing tumor growth for 21 days prior to treatment initiation (Extended Data Fig. 9a). At day 21, luciferase signals in the lung approached saturation levels (Extended Data Fig. 9b, c). While p53 R248Q-targeting LNPs did not reduce lung tumor burden in this advanced setting, they delayed metastasis formation compared to control treatments (Extended Data Fig. 9c). These results suggest that nucleic acids encoding Cas12a2 and suitable gRNAs could be co-delivered to target cancer-specific transcripts for tumor suppression.

## Discussion

The results reported here establish Cas12a2 RNP-mediated chromatin shredding as an effective approach to selectively eliminate cancer cells. We show that Cas12a2 cleaves eukaryotic chromatin *in trans* upon recognizing specific mRNA transcripts, triggering DNA damage responses and cell death in mammalian cells (Fig. 5g). This mechanism enabled us to target cancer cells with elevated *CCNE1* or *c-MYC* oncogene expression, *EGFR* in-frame deletion mutations and several *TP53* point mutations with high specificity.

The significance of these findings for cancer treatment could be profound, particularly for cancers with undruggable mutations such as *TP53* mutations. Instead of losing TP53 function through genetic deletion, cancer cells more commonly preserve clonal mutant copies (Extended Data Fig. 4a) that confer selective advantages during tumor evolution^6,7,26,34–36^. Consequently, nearly all tumor cells driven by *TP53* mutations retain mutant *TP53* transcript expression. By targeting these ubiquitous mutant transcripts, Cas12a2 could be impervious to tumor heterogeneity—a major challenge for drug resistance. Additionally, Cas12a2 can process its own CRISPR array^3^, enabling the potential for multiplexed targeting of multiple transcripts simultaneously. No approved methods exist to directly target *TP53* mutations, leaving a critical gap given p53’s importance in cancer. As the first approach to precisely target specific *TP53* mutations, our work paves the way for a new class of precision therapies using RNA-guided CRISPR nucleases.

## Methods

### Cell culture conditions

HEK293, RPE1 and U2OS cell lines were cultured in DMEM supplemented with 10% fetal bovine serum at 37 °C and 5% CO_2_. PC9 cells were cultured in RPMI1640 supplemented with 10% fetal bovine serum and 4mM total Glutamine at 37 °C and 5% CO_2_. Culture media were also supplemented with 1% penicillin/streptomycin.

### Cell lines

PC9 cells were obtained from Sigma (90071810). RPE1 p21-GFP and RPE1 p53 R248Q FUCCI cells were kindly gifted by John Diffley. U2OS TetON CycE was previously described in (Zeng et al., 2023) and was kindly gifted by John Diffley. HEKGFP High, HEKGFP Mid, HEKGFP Low, U2OS TetOn CycE LentiCas12a2, RPE1 p21-GFP LentiCas12a2, PC9 LentiCas12a2, RPE1 p53 R248Q LentiCas12a2, RPE1 p53 R175H LentiCas12a2, RPE1 p53 R280K LentiCas12a2, RPE1 p53 E285K LentiCas12a2 and RPE1 p53 R248Q FUCCI LentiCas12a2 were generated in this study.

Parental hTERT-RPE1 cells had puromycin resistance due to the presence of a puromycin resistance gene (PuroR) on the hTERT plasmid used for cell line immortalization. PuroR was knocked out in RPE1 cells using CRISPR-Cas9 with a gRNA sequence 5’ GCAACCTCCCCTTCTACGAG 3’. RPE1 single cell colonies were selected and loss of puromycin resistance was validated with 0.5 μg/ml puromycin. These RPE1 cells without PuroR were used for downstream cell line generation. RPE1 p21-GFP was generated with CRISPR-Cas9 knock-in as previously described^37^. For generating RPE1 p53 mutant cell lines, PLX313 plasmids carrying mutant *TP53* coding sequences and a Neomycin resistance gene (gifts from John Diffley) were used to make lentiviruses to transduce RPE1 cells at a MOI of 0.2. Stable cell lines were selected using 800 µg/ml G418. For generating LentiCas12a2 cells, plasmids carrying the SuCas12a2 coding sequence, tagged with a nucleoplasmin NLS at the N-terminus, followed by P2A-PuroR or P2A-TagBFP, were used to make lentiviruses as previously described^38^. After lentiviral transduction, stable lentiCas12a2 cell lines were selected using 2 µg/ml puromycin or confirmed with TagBFP expression.

PC9 cells expressing zsGreen-2A-Fluc (Firefly luciferase) were generated by lentiviral transduction. To introduce the Ai9 Cre-dependent tdTomato expression reporter into cancer cell lines, we made a Ai9 donor plasmid suitable for *in vitro* cell integration by adding inverted terminal repeats (ITRs) flanking the Ai9 expression cassette and puromycin resistance gene. The donor plasmid was then co-transfected with a Sleeping Beauty transposase-expressing plasmid pPGK-SB13 (Gift from Narita lab, Addgene 236078) into cells by lipofectamine 3000. Stable PC9 Ai9 cells were selected with 0.5 μg/ml puromycin.

For generating HEKGFP High, Mid and Low cells, single cell colonies were selected after transducing HEK293 cells with lentiviruses to express EGFP under a CMV promoter.

### Nucleic acid *trans*-cleavage assays

Reaction mixes contained 250 nM SuCas12a2, 300 nM gRNA, 250 nM target ssRNA, and 100 nM FAM-labeled collateral DNA or RNA substrate in 1x NEB3.1 buffer. For evaluating the effect of Mg²⁺ concentration, SuCas12a2, gRNA, and target ssRNA were pre-incubated at 37°C for 10 minutes in 1× NEB3.1 buffer (without MgCl₂), then combined with the FAM-labeled collateral substrate and different concentrations of MgCl₂ to initiate cleavage. Reactions were quenched at indicated timepoints by rigorous mixing with 10 µl phenol-chloroform; 4 µl of the aqueous layer after spinning was mixed with 4 µl 50% glycerol and resolved on 12.5% acrylamide Urea-Page denaturing gels, with cleavage products visualized on a Bio-Rad imager using the fluorescein channel.

### Chromatin *trans*-cleavage assays

The chromatinized plasmid was made with yeast histones using a 10,577 bp plasmid (pGCL42), containing 8,645 bp yeast sequence surrounding ARS1, kindly provided by Giselle Lee and John Diffley. 50 μg of purified histone octamers was mixed with 50 μg of pGCL42 to a final volume of 200 μL in buffer A (25 mM HEPES-KOH pH 7.6, 1 mM EDTA) containing 1 M NaCl. The histone-DNA mix was loaded into a D-Tube^TM^ Dialyser Mini (Merck Millipore) and dialysed at 4°C against 0.5 L buffer A of decreasing salt concentrations (1 M NaCl for 3 h, 0.75 M NaCl overnight, 0.5 M NaCl for 5 h, 0.0025 M NaCl overnight). After the final dialysis step, the dialysate is applied to a 5 ml 10%–40% v/v glycerol gradient (buffer A containing 0.0025 M NaCl) in a 13 × 51 mm tube (Beckman Coulter). Peak fractions containing reconstituted chromatin were pooled and assessed by micrococcal nuclease digestion. Chromatin was dialyzed against a storage buffer (25 mM HEPES-KOH pH 7.6, 2.5 mM NaCl, 0.1 mM EDTA) and stored at 4°C.

Chromatin cleavage reactions were performed in 10 µl volumes at 37°C in 1× NEB3.1 buffer. The mix contained 14 nM SuCas12a2, 14 nM gRNA, 25 nM target ssRNA, and with 1 nM of a naked DNA plasmid or the chromatinized plasmid. Reactions were initiated by pre-incubating SuCas12a2, crRNA, and target ssRNA at 37°C for 15 minutes to form the RNP-target complex, followed by the addition of either plasmid DNA or chromatin in a master stock. 10 µl reactions were taken out and quenched with 10 µl phenol-chloroform at indicated time points. The aqueous layer of quenched reactions was taken out and mixed with an equal volume of 50% glycerol before being analyzed on 1% agarose gels. For MNase control reactions, 0.04 U MNase was added to either 1 nM of the naked DNA plasmid or the chromatinized plasmid in 1× NEB3.1 buffer supplemented with 5 mM CaCl_2_ at 37 °C.

To assess Cas12a2’s ability to degrade mammalian chromatin, 200,000 HEK293T cells were harvested, washed once with PBS, and lysed in 100 µl CSK buffer (10 mM HEPES-KOH pH 7.9, 400 mM NaCl, 1 mM MgCl₂, 0.2% Triton X-100, 1 mM DTT, 100× protease-phosphatase inhibitor, 100 µg/ml BSA) at 4°C for 10 minutes. Then another 100 µl CSK buffer without NaCl was added to the lysed cells. Pre-incubated RNP-target RNA complex were prepared by incubating 250 nM SuCas12a2, 300 nM targeting gRNA or non-targeting gRNA, and 500 nM target ssRNA at 37°C for 10 minutes in 1× NEB buffer in 20 µ before adding to 200 µl nuclear extract (final concentrations: 250 nM SuCas12a2, 300 nM gRNA, 500 nM ssRNA). Reactions were incubated at 37°C, with aliquots taken at indicated time points, quenched with 300 nM NTI (From Takara NucleoSpin Gel and PCR Clean-Up Kit, 740609) buffer, and purified using the Takara kit, eluted in 15 µl elution buffer. Purified reaction products were visualized on 1% agarose gels.

### Nucleofection

A Cas12a2 RNP complex was prepared by combining Cas12a2 and gRNA (1:1 molar ratio) in 1× PBS (final RNP volume 5 µl) and incubated at room temperature for 10 minutes. 5.0 × 10⁵ HEK293 cells were centrifuged at 250 × g for 3 minutes, washed with 1× PBS, and resuspended in 20 µl nucleofection mix (16.4 µl SF Cell Line Solution, 3.6 µl Supplement 1; Lonza SF Cell Line 4D-Nucleofector X Kit) per condition before adding 5 µl RNP. 25 µl of the mixture was then transferred to a 16-well cuvette, nucleofected using the Lonza 4D-Nucleofector (program CA-189). Cells were then resuspended and seeded in 2 ml media in 6-wells. Incucyte (Sartorius) live-cell imaging was started 4 h post-seeding to monitor cell growth.

### Cas12a2 targeting in mammalian cells

For cells stably expressing Cas12a2, cells were seeded at ∼5% confluence (∼25,000 for RPE1, ∼20,000 for PC9, ∼40,000 for HEK293, ∼30,000 for U2OS) in 1 mL media in 24-well plates and reverse transfected with gRNAs. Transfection mixtures were prepared by combining Mix A (2 µL Lipofectamine RNAiMAX, 50 µL Opti-MEM) with Mix B (660 ng gRNA, 50 µL Opti-MEM), incubating for 30 min, before adding to each well. For co-transfecting Cas12a2 mRNA and gRNA, transfection mixtures were prepared by combining Mix A (1.5 µL Lipofectamine MessengerMAX, 50 µL Opti-MEM) with Mix B (150 ng Cas12a2 mRNA, 130 ng gRNA, 50 µL Opti-MEM), incubating for 30 min, before adding to each well. For LNP delivery (see LNP formulation below), 150 ng mRNA and 150 ng gRNA co-packaged in LNPs were added to each 24-well. Incucyte (Sartorius) live-cell imaging were started 4 h post-seeding to monitor cell growth.

### Live cell imaging analysis

Incutyte built-in AI confluence analysis and fluorescence analysis were used to measure cell confluence and GFP signal. For counting cell numbers, phase contrast images exported from Incucyte were segmented after Ilastik pixel classification training to identify individual cells. Segmented images were then analyzed using FIJI ImageJ to count cell numbers.

### *In vitro* transcription

Cas12a2 DNA templates were amplified via Polymerase Chain Reaction (PCR) using Q5 High-Fidelity DNA Polymerase (New England Biolabs) and purified by treatment with 0.8 U Proteinase K and 1/5 volume of 10% SDS at 37 °C for 1 h, followed by heat inactivation at 95 °C for 10 min. DNA was extracted by 1 volume of phenol–chloroform–isoamyl alcohol. After centrifugation at maximum speed, the aqueous phase was recovered, mixed with 1/10 volume of 3 M sodium acetate (pH 5.2) and 2 volumes of cold 100% ethanol, and precipitated at −20 °C overnight. DNA was pelleted by centrifugation (≥10 min, 4 °C), washed twice with ice-cold 70% ethanol, air-dried, and resuspended in 20–100 µL RNase-free water. SUPER RNase inhibitor (1 µL; Thermo Fisher Scientific) was added to each sample. All steps were carried out in an RNase-free PCR workstation.

RNA was synthesized using the HiScribe T7 High Yield RNA Synthesis Kit (New England Biolabs). Reactions were mixed at room temperature with ATP, GTP, CTP and pseudouridine-5′-triphosphate included at equimolar concentrations (10 mM final concentration each). Co-transcriptional capping was performed with CleanCap AG (TriLink, 10 mM final). Reactions were incubated at 37 °C for 4 h, followed by treatment with 1 µL Turbo DNase (Thermo Fisher Scientific) at 37 °C for 15 min. RNA was then purified by lithium chloride precipitation. The reaction mix was mixed with 0.5 volumes of 7.5 M LiCl and incubated at −20 °C for at least 1 h. Precipitated RNA was collected by centrifugation (15–30 min, 4 °C), washed with ice-cold 70% ethanol, centrifuged, and resuspended in RNase-free water. RNA concentration was measured by Nanodrop and RNA integrity was assessed using denaturing formaldehyde agarose gels.

### Development of *in vivo* liver tumor models

MYC liver tumor models were generated following previously published protocols^30,39^. Each female FVB/NJ mouse (Jackson Laboratory, Strain no. 001800) at 6-8 weeks of age was injected with 2 ml sterile PBS containing 10 μg pT3-EF1a-cMYC (Gift from Chen lab, Addgene 92046) and 1 μg pPGK-SB13 (Gift from Narita lab, Addgene 236078) plasmids via hydrodynamic tail vein injection (HDTVi) in 3 seconds. All animal handling, care, treatment and euthanasia procedures were performed in accordance with guidelines established by the relevant Institutional Animal Care and Use Committee (IACUC). Body and liver weights were measured at the time of sacrifice to calculate liver-to-body weight ratios. Tumor nodules and whole liver were annotated manually as ROIs by FIJI, and areas of ROIs were measured. The percentage of whole liver area covered by tumors is calculated as follows: (sum of areas covered all tumors)/(whole liver area).

### Development of *in vivo* lung tumor model

PC9 cells were injected intravenously into the tail vein of NOD.Cg-*Prkdc^scid^ Il2rg^tm1Wjl^*/SzJ (NSG) mice (Jackson Laboratory, Strain no. 005557) in 0.1-0.2 mL PBS to generate orthotopic tumors of NSCLC. For bioluminescence measurement of tumor burden, mice were injected intraperitoneally with 150 mg/kg D-Luciferin 10 min before being imaged using an IVIS spectrum imager. Bioluminescence images were analyzed on Living Image software. For delivery efficiency assessment, lung tissues were harvested, dissociated into single cells by digestion with collagenase and passing through 40 μm filters, before analysis by flow cytometry. All animal handling, care, treatment and euthanasia procedures were performed in accordance with guidelines established by the relevant Institutional Animal Care and Use Committee (IACUC).

### Lipid nanoparticle formulation

4A3-SC8 (Catalog no. HY-148559) and 1,2-Dioleoyl-3-dimethylammonium-propane (DODAP, Catalog no. HY-130751) were purchased from MedChemExpress. 1,2-dioleoyl-sn-glycero-3-phosphoethanolamine (DOPE, Catalog no. 850725) and 1,2-Dimyristoyl-rac-glycero-3-methylpolyoxyethylen (DMG-PEG₂K, Catalog no.880151) were purchased from Avanti Polar Lipids. Cholesterol (Catalog no. C8667) was purchased from Sigma. 1,2-dioleoyl-3-trimethylammonium-propane (DOTAP, Catalog no. D6182) was purchased from Sigma. For liver tumor treatment experiments, three doses of 10% DOTAP SORT (Selective Organ Targeting) LNP and three doses of 20% DODAP SORT LNP were used and formulated based on previously published protocols^40^. For lung tumor treatment experiments, the 40% DOTAP SORT LNP composition was used and formulated. The molar ratios for 4A3-SC8, DOPE, cholesterol. DME-PEG and DOTAP in 10% DOTAP and 40% DOTAP LNPs are 14.3:14.3:28.5:2.9:40 and 21.5:21.4:42.8:4.3:10 respectively. The molar ratio for 4A3-SC8, DOPE, cholesterol. DME-PEG and DODAP in 20% DODAP is 19.1:19:38.1:3.8:20. The total lipid to RNA weight ratio is 40:1 for 10% DOTAP and 40% DOTAP LNPs, and 20:1 for 20% DODAP LNPs. Lipids were dissolved in ethanol and mixed by microfluidics (NanoAssemblr Ignite) with RNA diluted in 10 mM citrate buffer (pH 4.0) at a lipid-to-RNA volume ratio of 1:3. The RNA payload consisted of mRNA and guide RNA combined at a 1:1 mass ratio. LNPs were then dialyzed (PurA-Lyzer Midi Dialysis Kits, WMCO 3.5 kDa, Catalog no. PURX35100) over night before being concentrated by centrifugation (Amicon Centrifugal Filter 30 kDa) at 2,000 g. Dynamic light scattering (DLS) measurements of LNPs were performed using Benano 180 Zeta Pro (Suzhou Benano Nanotech Co., Ltd., Cat. No. DLS180-Pro). In the *in vivo* anti-tumor tests, LNPs were dosed at 2 mg total RNA per kg body weight dissolved in 0.1-0.2 mL PBS via the lateral tail vein.

### RNA Single Molecule Fluorescence in Situ Hybridization (smFISH)

Cells were plated on coverslips, fixed with 4% paraformaldehyde (PFA) in PBS at room temperature (RT) for 15 minutes, washed three times with 1× PBS, permeabilized with 0.5% Triton X-100 in PBS for 10 minutes, and washed again three times with 1× PBS. Cells were then incubated in 30% formamide wash buffer at RT for at least 5 minutes, followed by hybridization with 50 µl of 3× hybridization buffer (30% vol/vol formamide, 0.1% wt/vol yeast tRNA, 0.1% vol/vol murine RNase inhibitor, 10% vol/vol dextran sulfate in 2× SSC) containing RNA smFISH probes at 37°C overnight. Samples were then washed twice with 30% formamide wash buffer at 37°C for 30 minutes each, followed by two washes with 2× SSC, and either stored in 2× SSC with RNase inhibitor or immediately processed for readout staining by incubating with 10% ethylene carbonate (EC) hybridization buffer (10% vol/vol EC in 2× SSC) containing 3 nM readout probe at RT for 15 minutes, washing twice with 2× SSC, and imaging on a confocal microscope with 63x or 100x oil immersion lens.

### Antibodies

Immunoblotting was performed using the following antibodies diluted in TBS buffer supplemented with 0.1% Tween 20 and 5% milk powder or 3% BSA: p-KAP1 (1:1000, Bethyl Laboratories, A300-767A), p-H2A.X S139 (1:1000, Millipore, 05-636), GAPDH (1:1000, Santa Cruz, sc-365062), anti-Mouse HRP(1:5000, Invitrogen, 31430), anti-Rabbit HRP (1:5000, Invitrogen, 65-6120), anti-Rabbit IRDye800 (1:5000, Licor, 926-32211) and anti-Mouse IRDye680 (1:5000, Licor, 926-68070).

## Acknowledgments

We thank Yehui Sun, Stephen Moore, Daniel J. Siegwart, Xin Chen and Chase L. Biesel for technical advice or insightful discussions. J.A.D. is an investigator of the Howard Hughes Medical Institute (HHMI) and this research is supported by the CRISPR Cures for Cancer fund and Gladstone Institutes. J.A.D. also receives support from NIH/NIAID (U54AI170792, UH3AI150552 and U01AI142817), NIH/NINDS (U19NS132303), NIH/NHLBI (R21HL173710), NSF (2334028), DOE (DE-AC02-05CH11231, 2553571 and B656358); Lawrence Livermore National Laboratory, Apple Tree Partners (24180), UCB-Hampton University Summer Program, Mr. Li Ka Shing, Koret-Berkeley-TAU, Emerson Collective and the Innovative Genomics Institute (IGI). The Gladstone Institutes acknowledges the generous support of the James B. Pendleton Charitable Trust. HHMI has covered open publication access charges. Y.L. acknowledges support from NIGMS (R35GM150941).

## Author Contributions

Conceptualization: J.Z. and J.A.D.

Proof-of-concept demonstration of mammalian cell killing: J.T. and Y.L.

Design and conceptualization of *in vivo* studies: J.Z., H.C., H.H., N.M., J.A.D., and A.A.

Materials and Methodology: J.Z., Z.C., H.C., H.H., J.T., K.T.C., W.N., C.X., D.R., M.H.K., G.L., J.F.X.D., Y.S., Y.M., L.Q., N.M.K., R.N.J., Y.L., N.M., and A.A.

Biochemistry experiments: J.Z., K.T.C., and J.T

Mammalian cell killing experiments: J.Z., Z.C. J.T

*In vitro* RNA transcription: W.N., D.R., J.Z., and Z.C.

*In vivo* tests: H.C., J.Z., H.H., and Z.C.

RNA smFISH: C.X.

Writing—original draft: J.Z., Z.C. and J.A.D.

Writing—review and editing: all of the authors.

Visualization: J.Z.

Supervision: J.A.D.

## Extended Data Figures and Legends

**Extended data fig 1.**
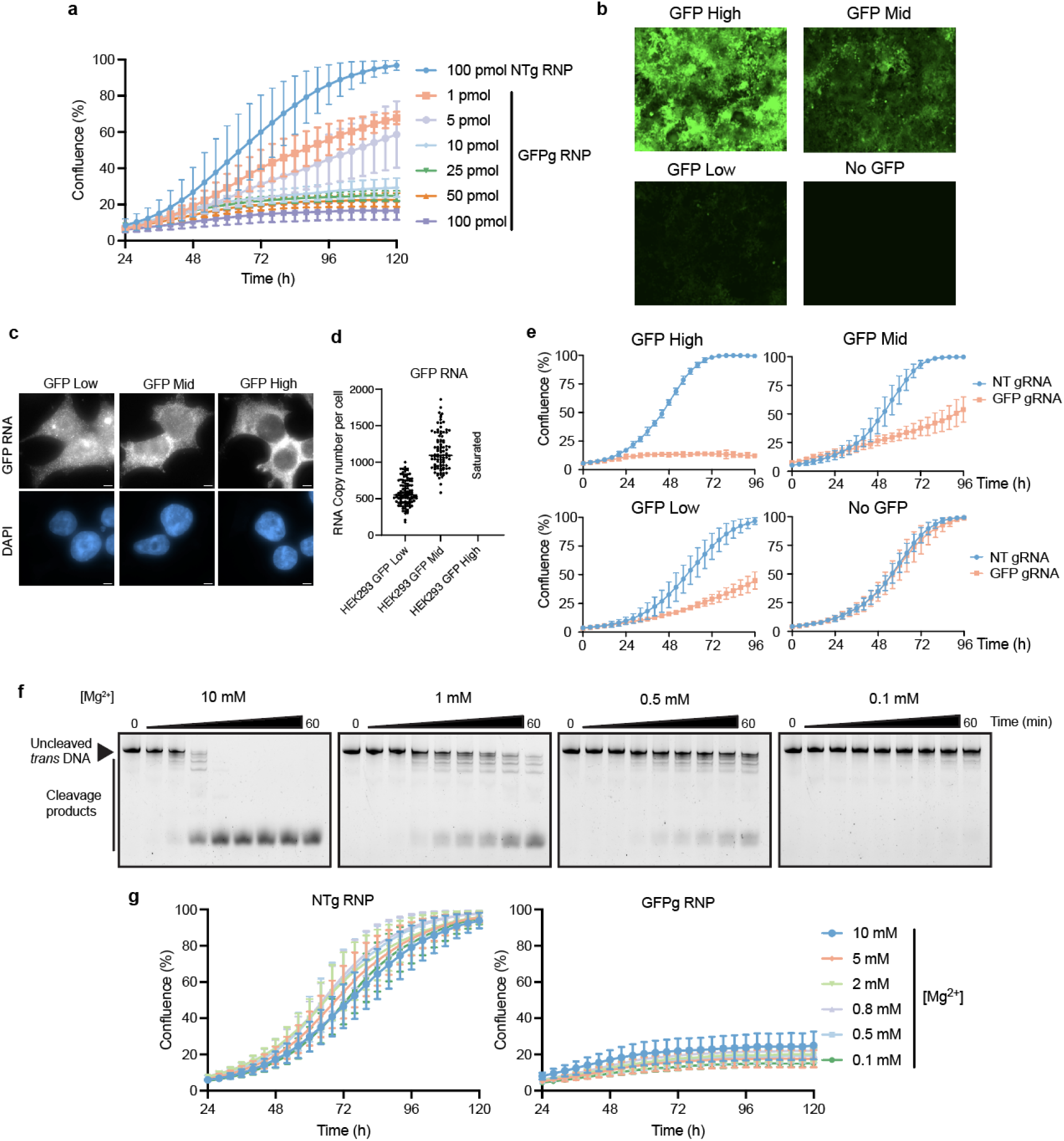
**a,** Growth curves of HEK293-GFP (mid-level expression in **b**) cells nucleofected with different concentrations of Cas12a2-GFP gRNA RNP. **b,** Green channel images of HEK293 cells expressing different levels of GFP (High, Mid, Low). **c,** Representative smFISH cell images. Scale bar, 5 µm. **d,** Quantification of RNA numbers from smFISH images. Bar indicates the median. **e,** Growth curves of HEKGFP cells in **b** nucleofected with 5pmol of Cas12a2-GFP gRNA RNP. **f,** Time-course *in vitro* DNA trans-cleavage analysis by Cas12a2 in different concentrations of Mg^2+^. **g,** Growth curves of HEK293-GFP (mid) cells nucleofected with Cas12a2-gRNA RNP pre-incubated in different concentrations of Mg^2+^.

**Extended data fig 2.**
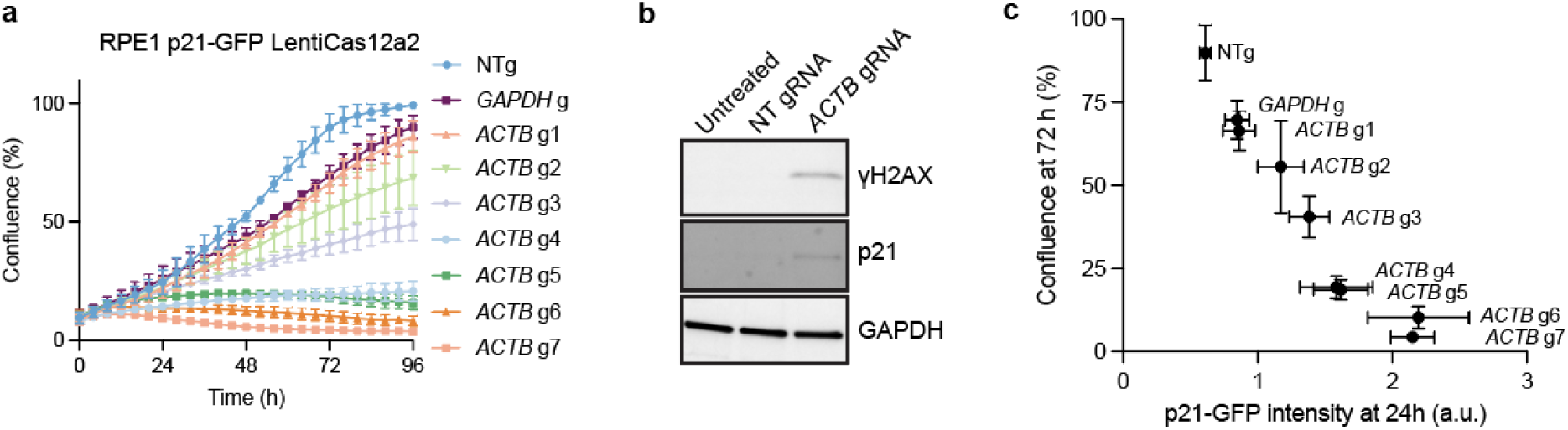
**a,** Growth curves of RPE1 p21-GFP LentiCas12a2 cells transfected with gRNAs targeting *ACTB* or *GAPDH.* **b,** Immunoblots showing expression of DNA damage markers in RPE1 p21-GFP LentiCas12a2 cells 48 h following transfecting *ACTB* g7. **c,** Correlation of cell confluence of RPE1 p21-GFP LentiCas12a2 cells in **a** with p21-GFP intensity.

**Extended data fig 3.**
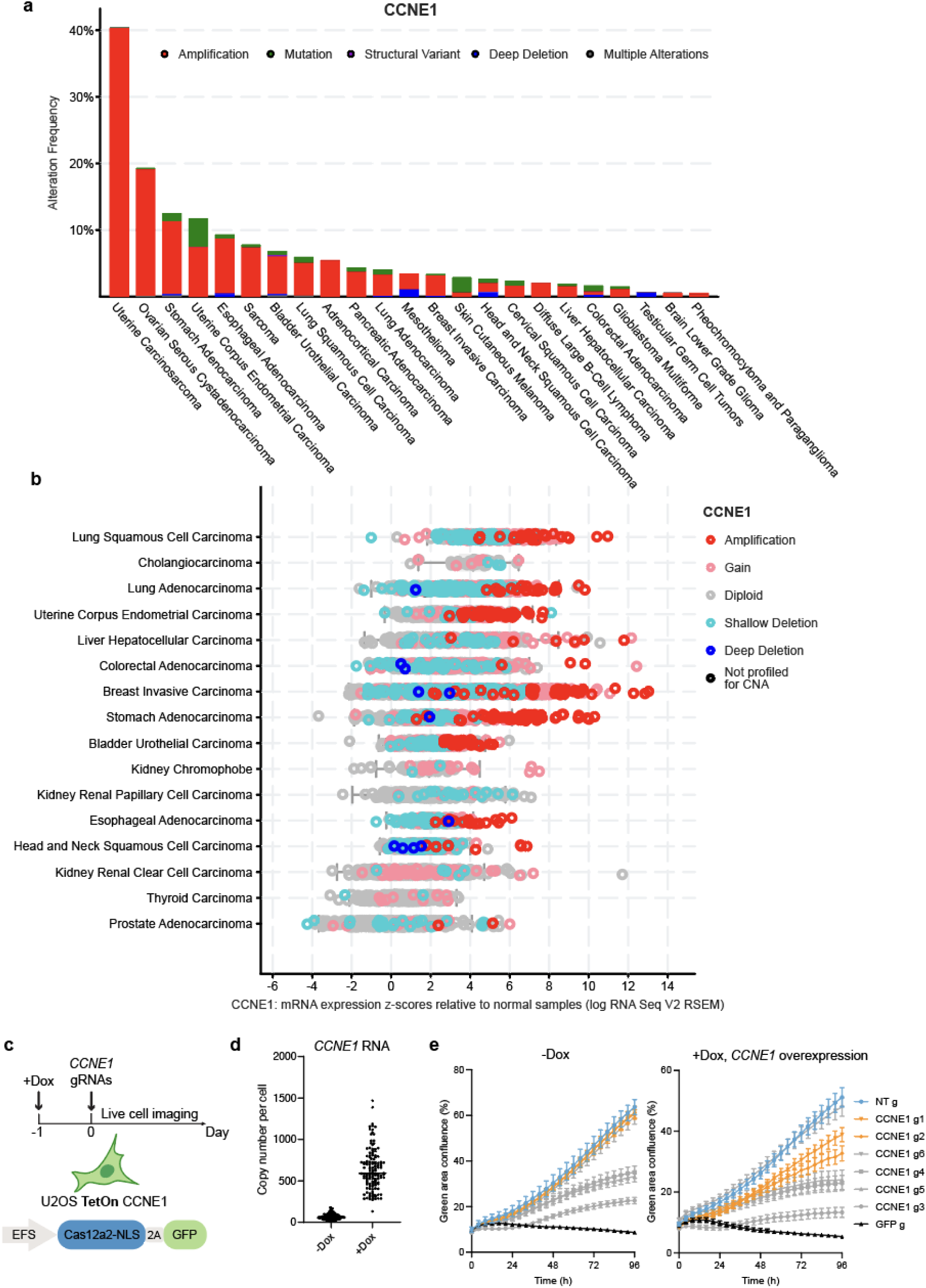
**a,** Prevalence of *CCNE1* alterations in different cancers from TCGA Pan-Cancer Atlas studies. **b,** *CCNE1* mRNA expression levels in different cancers from TCGA Pan-Cancer Atlas studies. **c,** Testing of the *CCNE1*-targeting gRNAs in U2OS cells expressing CCNE1 under a Dox-inducible promoter and stably expressing Cas12a2. **d,** Quantification of RNA levels. Bar indicates the median. **e,** Growth curves of U2OS cells in **c.**

**Extended data fig 4.**
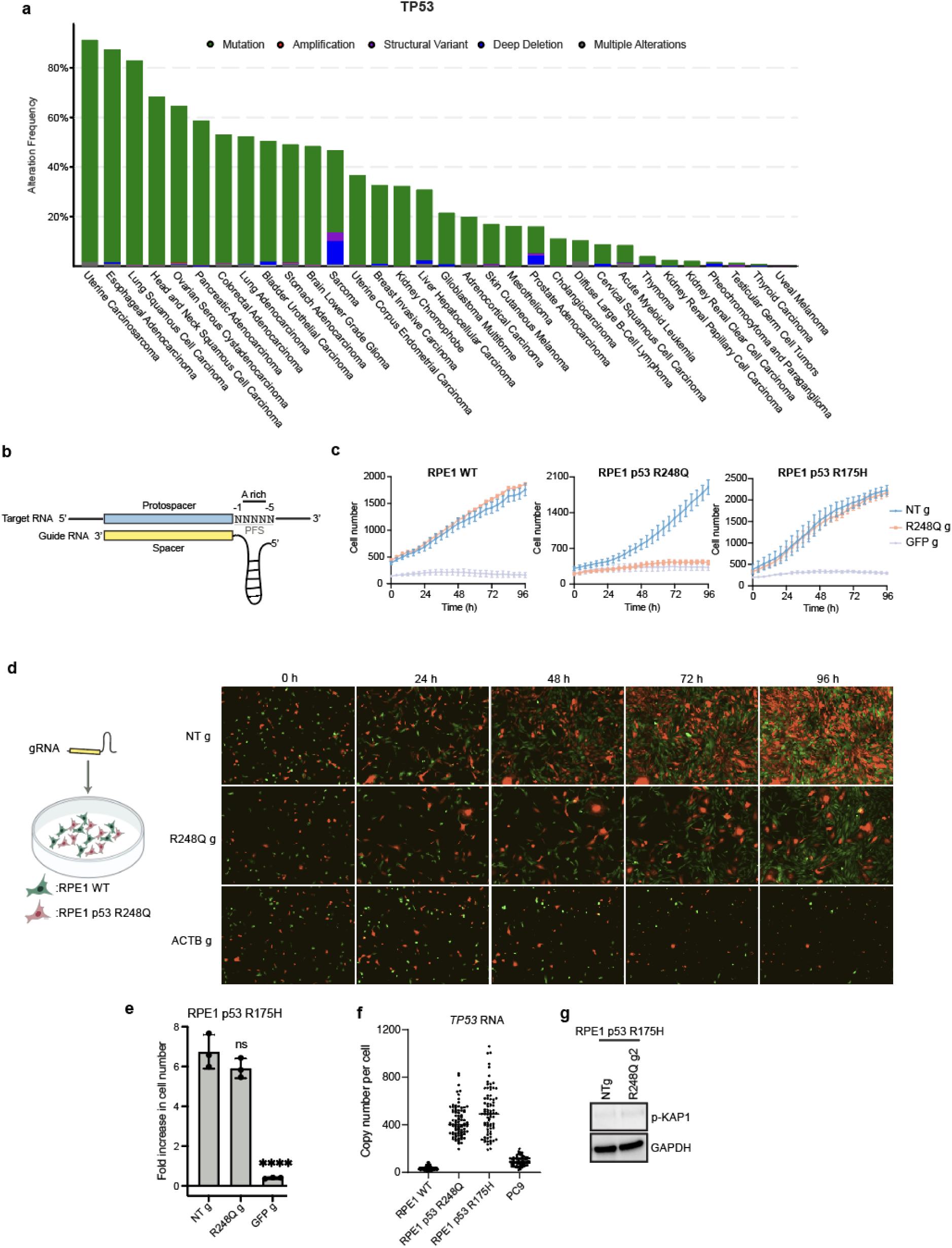
**a,** Prevalence of *TP53* alterations in different cancers from TCGA Pan-Cancer Atlas studies. **b,** Schematic showing the protospacer flanking site (PFS) for Cas12a2 targeting. **c,** Growth curves of RPE1 WT and p53 mutant cells following Cas12a2 targeting of p53 R248Q mRNA. **d**, Growth competition assay between RPE1 WT and R248Q cells, both stably expressing Cas12a2. Images of merged red and green channels are shown. **e,** Fold-increase in cell number over 96 h following Cas12a2 targeting of the p53 R248Q mutant transcript in RPE1 p53 R175H control cells. **f,** Quantification of RNA levels using smFISH. Bar indicates the median. **g,** Immunoblots probing expression of DNA damage marker phospho-KAP1 48 h following Cas12a2 targeting of the p53 R248Q mutant transcript in RPE1 p53 R175H control cells. Statistics: ****p<0.0001; ns, non-significant, one-way ANOVA with Dunnett’s test.

**Extended data fig 5.**
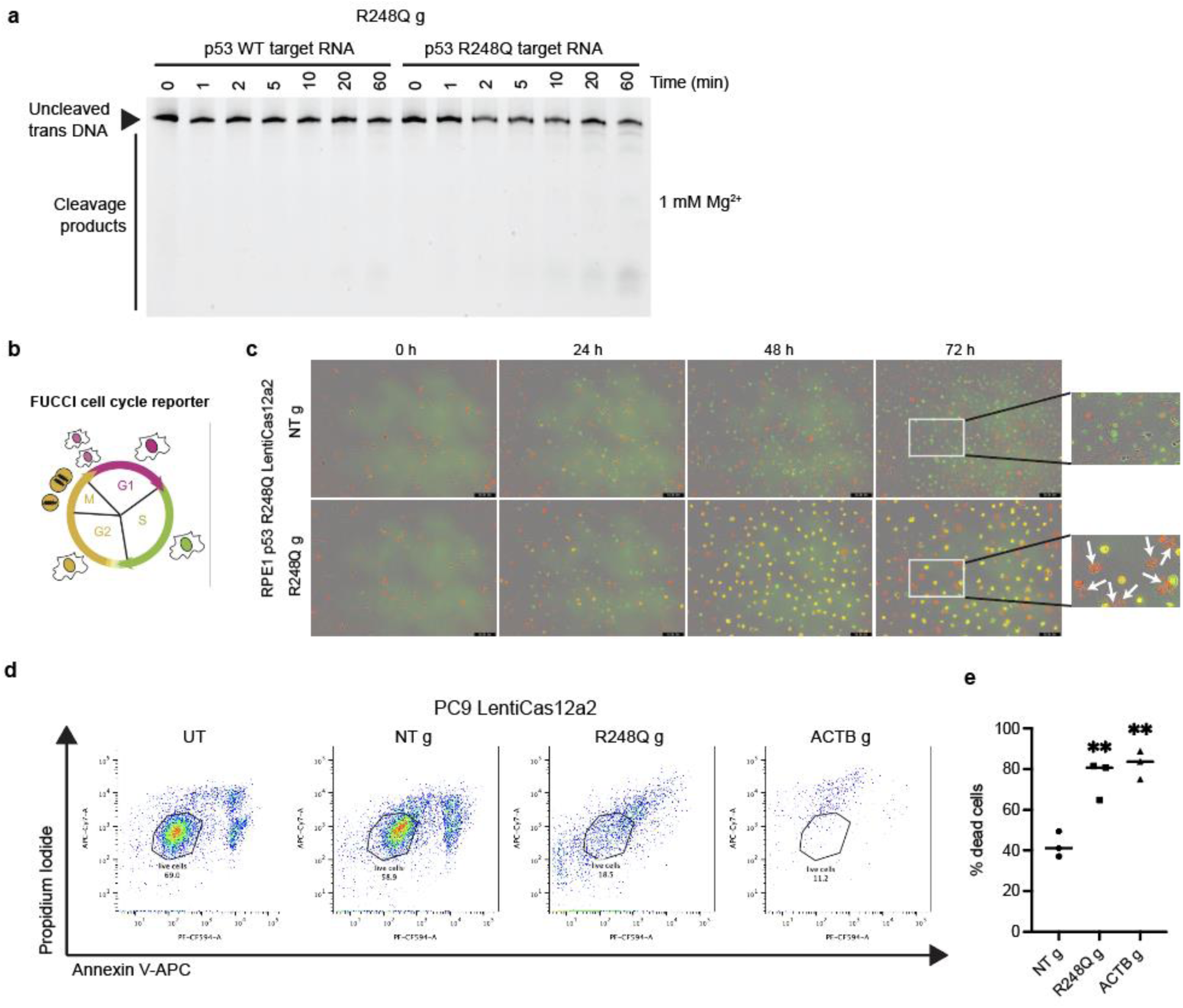
**a,** Time-course analysis of *in vitro trans*-cleavage of FAM-labelled dsDNA by Cas12a2 with R248Q g in the presence of p53 R248Q RNA fragment or p53 WT RNA fragment. **b,** Schematic of the FUCCI cell cycle reporter. **c,** Time-lapse images of RPE1 p53 R248Q cells expressing the FUCCI reporter following Cas12a2 targeting of the p53 R248Q mutant transcript. Merged green, red and phase contrast channels are shown. White arrows indicate cells with fragmented nuclei. **d,** FACS gating of dead cell populations in PC9 LentiCas12a2 cells 96 h following Cas12a2 targeting. **e,** Quantification of dead cell populations in PC9 LentiCas12a2 cells 96 h following Cas12a2 targeting. Statistics: **p<0.01, one-way ANOVA with Dunnett’s test.

**Extended data fig 6.**
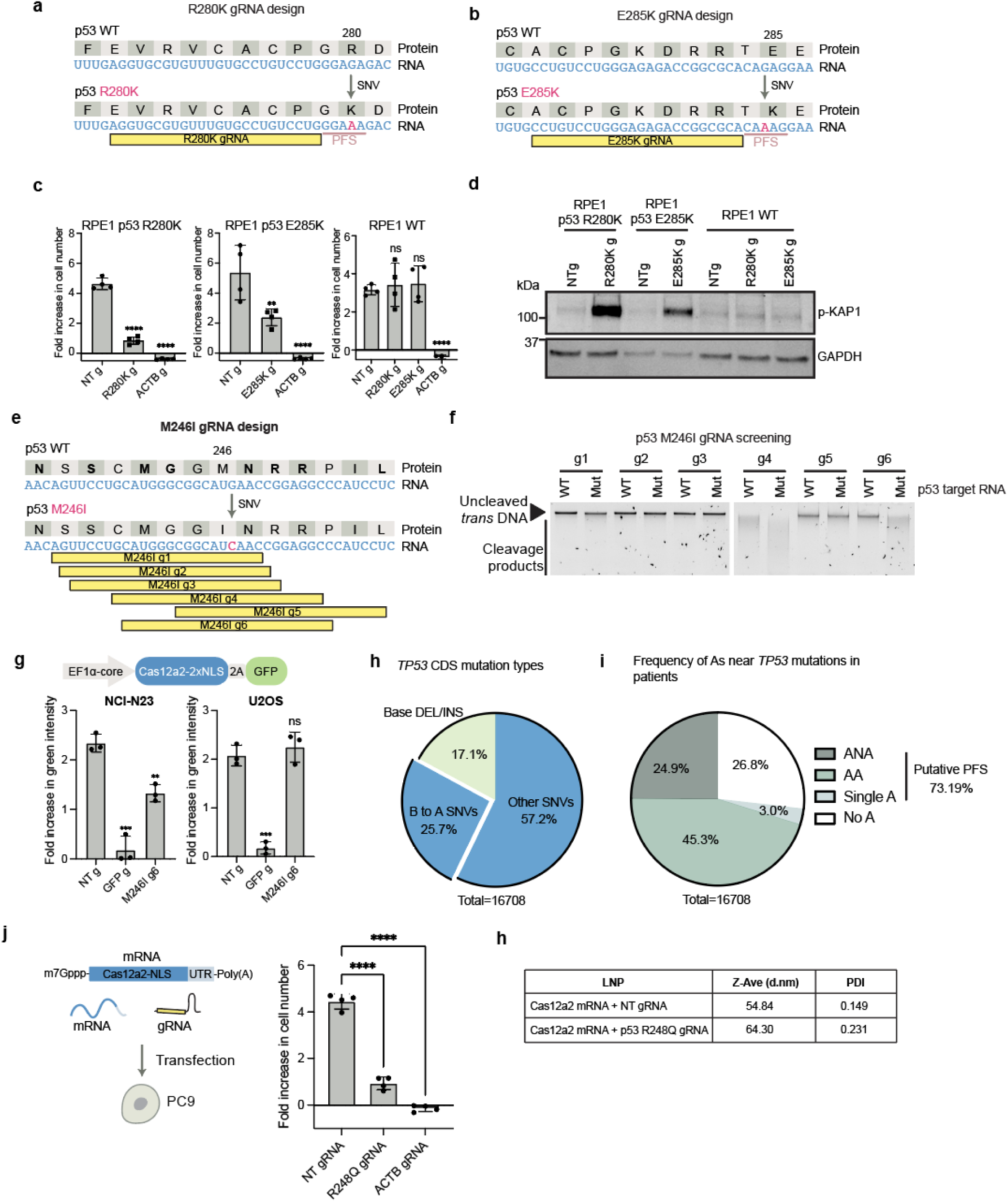
**a,** Design of a gRNA (R280K g) targeting the p53 R280K mutant transcript. **b,** Design of a gRNA (E285K g) targeting the p53 E285K mutant transcript. **c,** Fold-increase in cell number over 96 h following Cas12a2 targeting of p53 R280K or E285K mutant transcripts. **d,** Immunoblots showing expression of DNA damage marker phospho-KAP1 in RPE1 p53 R280K, RPE1 p53 E285K and RPE1 WT cells 48 h following Cas12a2 targeting of the mutant transcripts. **e,** Design of gRNAs (M246I g1-6) targeting the p53 M246I mutant transcript. **f,** *Trans*-cleavage of purified genomic DNA by Cas12a2 with M246I gRNAs in the presence of the mutant target RNA or WT target RNA. **g**, Fold-increase in green intensity over 96 h following Cas12a2 targeting of the p53 M246I mutant transcript in NCI-H23 cells (p53 M246I) and U2OS cells (p53 WT) expressing Cas12a2-2A-EGFP. **h,** Analysis of mutation types in the *TP53* coding sequence (CDS) from 16,708 tumor samples. Although Cas12a2 prefers an A-rich PFS (Extended Data Fig. 4b), a consensus PFS sequence could not be built^3^. However, we noticed that a single adenine nucleotide can sometimes serve as a PFS to activate Cas12a2 (Extended Data Fig. 6e, f), and more adenine bases between the -2 to -4 position at the 3’ end of the protospacer can lead to higher Cas12a2 activity (Fig. 4b-d and Extended Data Fig. 6a-d). 25.7% of *TP53* mutations in patients are B-to-A substitutions (where B is G, C, or T) (Extended Data Fig. 6h), which could serve as new or enhanced PFS for targeting, as demonstrated with the R248Q mutation (Fig. 4b-d). 17.1% of *TP53* mutations are base insertions or deletions (Extended Data Fig. 6h), offering opportunities for targeting similar to our approach with the *EGFR* in-frame deletion mutation (Fig. 3a-e). **i,** Frequency of the presence of A, AA or ANA within 24 nt at the 3’ end of *TP53* mutations from 16, 708 tumor samples. Tumor sample data in **c** and **d** are from TCGA Pan-Cancer Atlas studies and MSK-CHORD studies. **j,** Cas12a2 targeting by transfecting mRNA encoding Cas12a2 and gRNAs into PC9 cells. Growth curves and fold-increase in cell number over 96 h are shown on the right. **h,** Characterization of LNPs used for the *in vivo* lung tumor treatment test. Statistics: **p<0.01; ****p<0.0001; ns, non-significant, one-way ANOVA with Dunnett’s test.

**Extended data fig 7.**
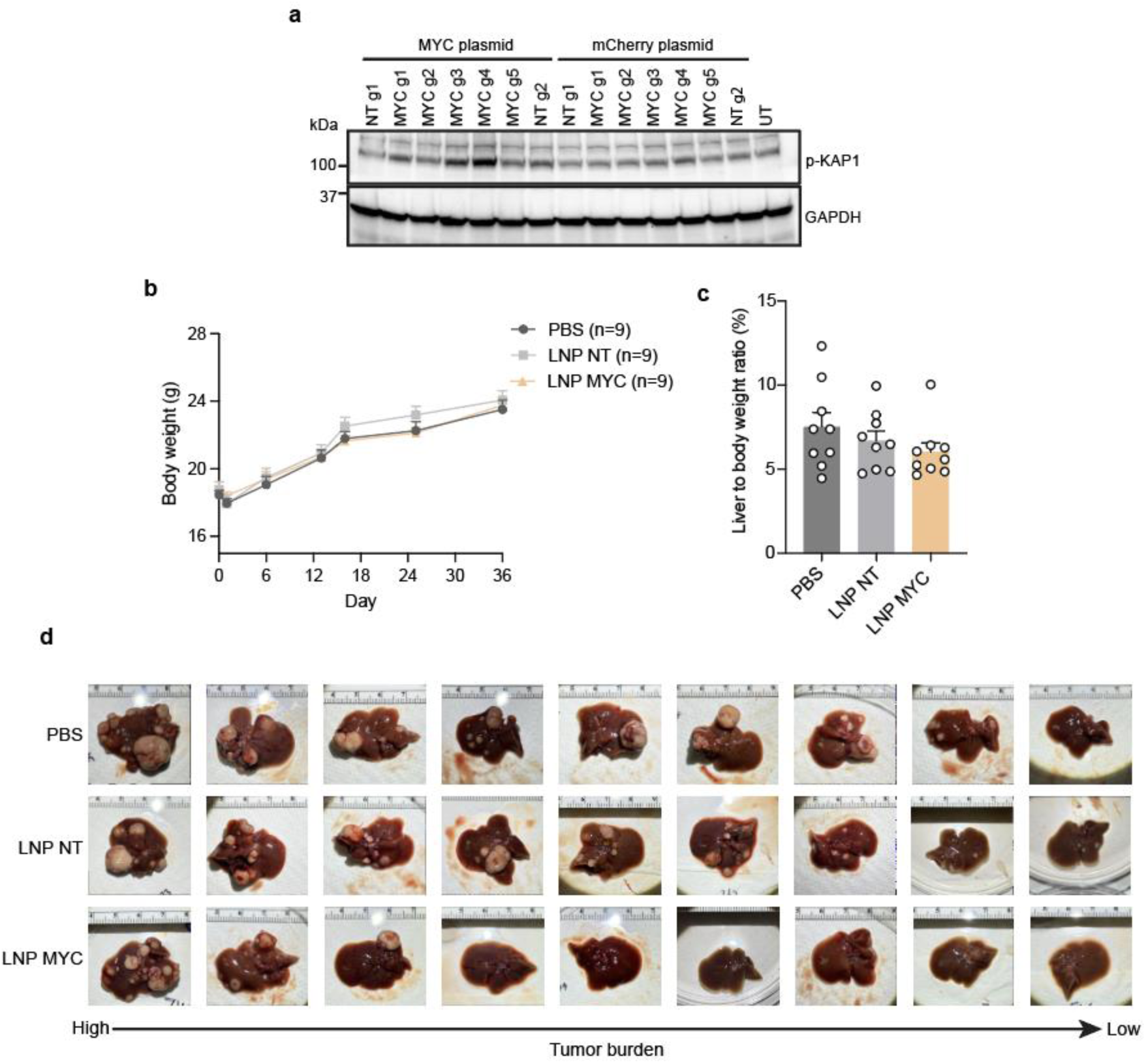
**a,** HEK293 cells expressing Cas12a2 were transfected with a MYC-encoding plasmid or mCherry-encoding control plasmid. 24 hours later gRNAs targeting MYC were transfected. Immunoblots probing expression of DNA damage marker phospho-KAP are shown. **b**, Changes in body weight of mice in Fig. 5c. **c**, Quantification of liver to body weight ratios of mice in Fig. 5c at the endpoint. **d,** Endpoint liver images of mice in Fig. 5c.

**Extended data fig 8.**
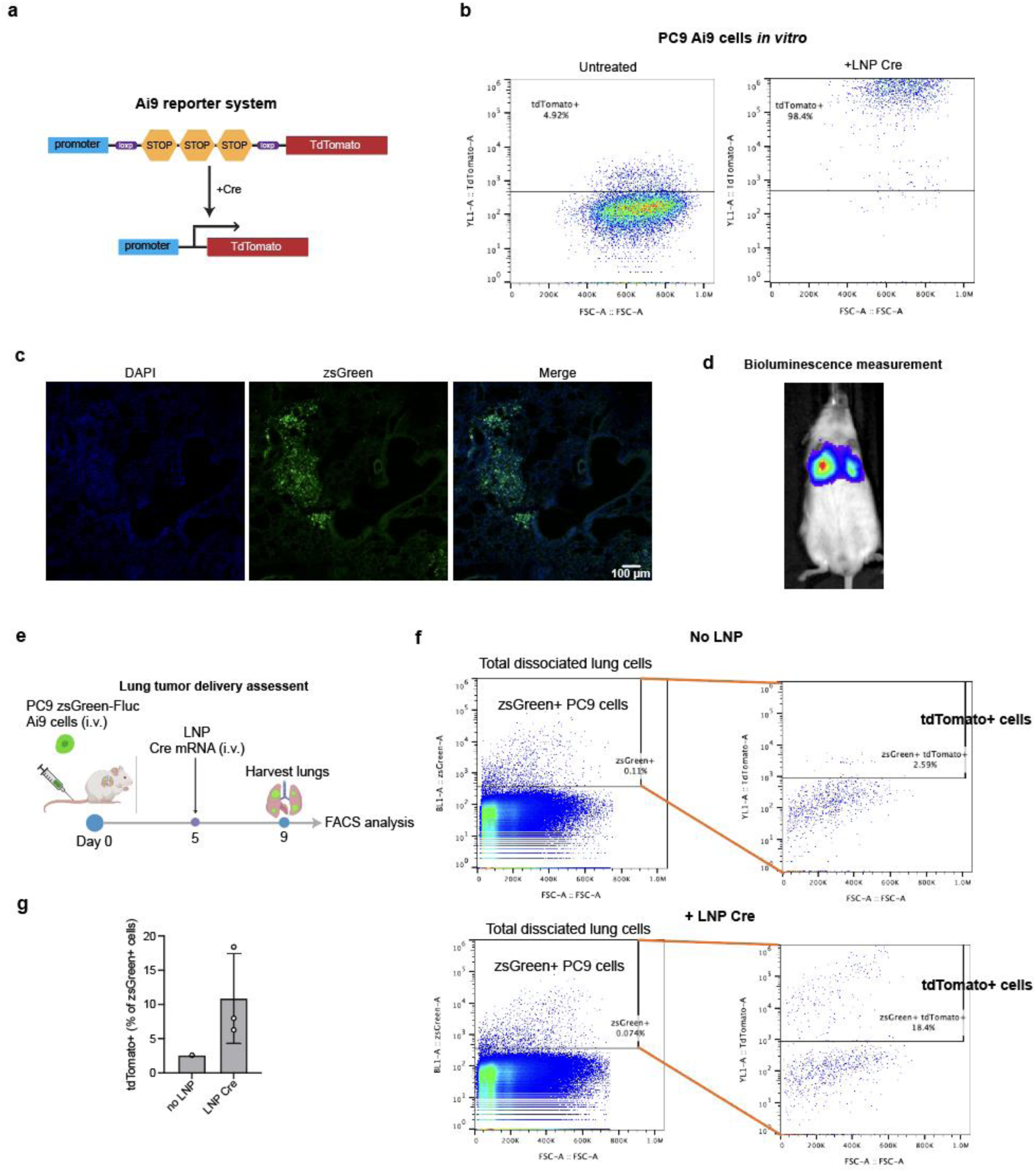
**a,** Schematic showing the Ai9 Cre-dependent tdTomato reporter system. The reporter construct was integrated into PC9 cells by the Sleeping Beauty transposase. Cells were selected by puromycin. **b,** Flow cytometry analysis showing tdTomato expression in PC9 Ai9 cells after treating them with LNPs delivering Cre mRNA *in vitro*. **c,** Fluorescent histological images of a mouse lung 53 days post engraftment with PC9 zsGreen-Fluc cells. **d,** IVIS image of a mouse injected with PC9 zsGreen-Fluc cells. **e,** Schematic illustrating delivery efficiency assessment of LNPs to PC9 lung tumors. Each mouse received 20 μg Cre mRNA delivered by LNP. **f and g,** Flow cytometry analysis of isolated PC9 zsGreen Ai9 cells from mouse lung tissues after LNP Cre mRNA delivery as shown in **e.** Quantification of the percentage of tdTomato+ cells is shown in **g** (n=3 mice).

**Extended data fig 9.**
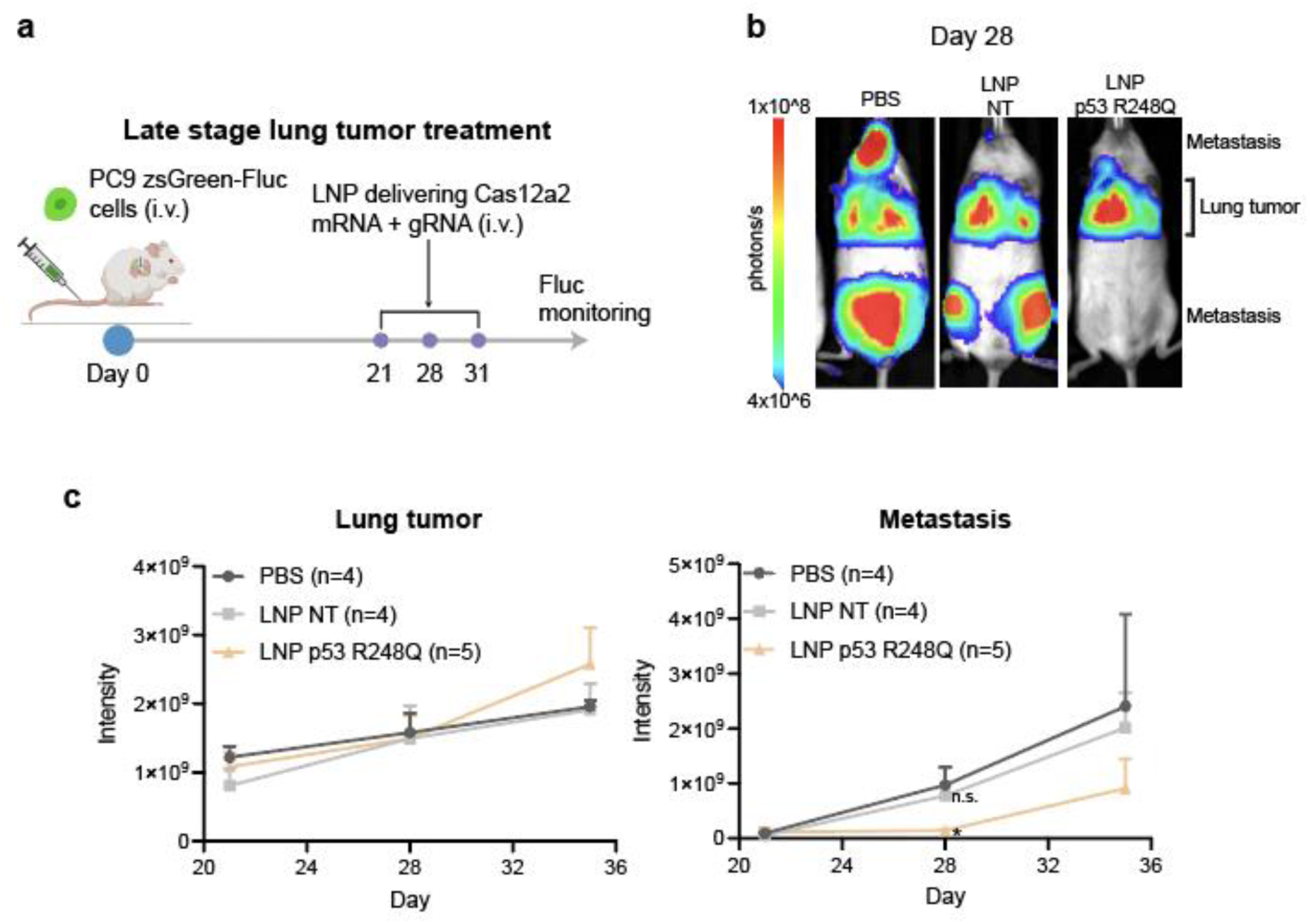
**a,** Schematic showing late-stage lung tumor treatment scheme. **b,** Representative IVIS images of mice bearing Fluc-expressing PC9 cells. Bioluminescence signals outside the lung are considered from metastasis. **c,** Quantification of tumor bioluminescence signals in the lung and outside the lung over time in treated mice in **a**.

## References

1. Jee, J. et al. Automated real-world data integration improves cancer outcome prediction. Nature 636, 728–736 (2024).

2. Zehir, A. et al. Mutational landscape of metastatic cancer revealed from prospective clinical sequencing of 10,000 patients. Nat. Med. 23, 703–713 (2017).

3. Dmytrenko, O. et al. Cas12a2 elicits abortive infection through RNA-triggered destruction of dsDNA. Nature 613, 588–594 (2023).

4. Bravo, J. P. K. et al. RNA targeting unleashes indiscriminate nuclease activity of CRISPR–Cas12a2. Nature 613, 582–587 (2023).

5. Hainaut, P. & Pfeifer, G. P. Somatic TP53 Mutations in the Era of Genome Sequencing. Cold Spring Harb. Perspect. Med. 6, a026179 (2016).

6. Gerstung, M. et al. The evolutionary history of 2,658 cancers. Nature 578, 122–128 (2020).

7. Consortium, T. I. P.-C. A. of W. G., et al. Pan-cancer analysis of whole genomes. Nature 578, 82–93 (2020).

8. Levine, A. J. & Oren, M. The first 30 years of p53: Growing ever more complex. Nature Reviews Cancer 9, 749–758 (2009).

9. Joerger, A. C. & Fersht, A. R. The Tumor Suppressor p53: From Structures to Drug Discovery. Cold Spring Harb. Perspect. Biol. 2, a000919 (2010).

10. Sabapathy, K. & Lane, D. P. Therapeutic targeting of p53: all mutants are equal, but some mutants are more equal than others. Nat. Rev. Clin. Oncol. 15, 13–30 (2018).

11. Abraham, C. G. & Espinosa, J. M. The Crusade against Mutant p53: Does the COMPASS Point to the Holy Grail? Cancer Cell 28, 407–408 (2015).

12. Duffy, M. J., Synnott, N. C., O’Grady, S. & Crown, J. Targeting p53 for the treatment of cancer. Semin. Cancer Biol. 79, 58–67 (2022).

13. Xiao, S. et al. Characterization of the generic mutant p53-rescue compounds in a broad range of assays. Cancer Cell 42, 325–327 (2024).

14. Johmura, Y. et al. Necessary and sufficient role for a mitosis skip in senescence induction. Molecular Cell 55, 73–84 (2014).

15. Krenning, L., Feringa, F. M., Shaltiel, I. A., vandenBerg, J. & Medema, R. H. Transient activation of p53 in G2 phase is sufficient to induce senescence. Molecular Cell 55, 59–72 (2014).

16. Zeng, J., Hills, S. A., Ozono, E. & Diffley, J. F. X. Cyclin E-induced replicative stress drives p53-dependent whole-genome duplication. Cell 186, 528–542.e14 (2023).

17. White, D. et al. The ATM Substrate KAP1 Controls DNA Repair in Heterochromatin: Regulation by HP1 Proteins and Serine 473/824 Phosphorylation. Mol. Cancer Res. 10, 401–414 (2012).

18. Romani, A. & Scarpa, A. Regulation of cell magnesium. Arch. Biochem. Biophys. 298, 1–12 (1992).

19. Eggers, A. R. et al. Rapid DNA unwinding accelerates genome editing by engineered CRISPR-Cas9. Cell 187, 3249–3261.e14 (2024).

20. Linzer, D. I. H. & Levine, A. J. Characterization of a 54K Dalton cellular SV40 tumor antigen present in SV40-transformed cells and uninfected embryonal carcinoma cells. Cell 17, 43–52 (1979).

21. Lane, D. P. & Crawford, L. V. T antigen is bound to a host protein in SY40-transformed cells. Nature 278, 261–263 (1979).

22. Etemadmoghadam, D. et al. Synthetic lethality between CCNE1 amplification and loss of BRCA1. Proc. Natl. Acad. Sci. 110, 19489–19494 (2013).

23. Bartkova, J. et al. DNA damage response as a candidate anti-cancer barrier in early human tumorigenesis. Nature 434, 864–870 (2005).

24. Mitsudomi, T. & Yatabe, Y. Mutations of the epidermal growth factor receptor gene and related genes as determinants of epidermal growth factor receptor tyrosine kinase inhibitors sensitivity in lung cancer. Cancer Sci. 98, 1817–1824 (2007).

25. Sharma, S. V., Bell, D. W., Settleman, J. & Haber, D. A. Epidermal growth factor receptor mutations in lung cancer. Nat. Rev. Cancer 7, 169–181 (2007).

26. Freed-Pastor, W. A. & Prives, C. Mutant p53: one name, many proteins. Genes & Development 26, 1268–1286 (2012).

27. Levine, A. J., Momand, J. & Finlay, C. A. The p53 tumour suppressor gene. Nature 351, 453–456 (1991).

28. Sakaue-Sawano, A. et al. Genetically Encoded Tools for Optical Dissection of the Mammalian Cell Cycle. Molecular Cell 68, 626–640.e5 (2017).

29. Sun, Y. et al. In vivo editing of lung stem cells for durable gene correction in mice. Science 384, 1196–1202 (2024).

30. Chow, E. K., Fan, L., Chen, X. & Bishop, J. M. Oncogene-specific formation of chemoresistant murine hepatic cancer stem cells. Hepatology 56, 1331–1341 (2012).

31. Zhao, S. et al. Acid-degradable lipid nanoparticles enhance the delivery of mRNA. Nat. Nanotechnol. 19, 1702–1711 (2024).

32. Chen, K. et al. Lung and liver editing by lipid nanoparticle delivery of a stable CRISPR–Cas9 ribonucleoprotein. Nat. Biotechnol. 1–13 (2024) doi:10.1038/s41587-024-02437-3.

33. Lee, Y. et al. Lung-targeted delivery of TRAIL and BAK mRNA by optimized lipid nanoparticles for in vivo lung metastasis. Chem. Eng. J. 522, 167379 (2025).

34. Olive, K. P. et al. Mutant p53 gain of function in two mouse models of Li-Fraumeni syndrome. Cell 119, 847–860 (2004).

35. Soussi, T. & Wiman, K. G. TP53: An oncogene in disguise. Cell Death and Differentiation 22, 1239–1249 (2015).

36. Muller, P. A. J. & Vousden, K. H. p53 mutations in cancer. Nat. Cell Biol. 15, 2–8 (2013).

37. Barr, A. R. et al. DNA damage during S-phase mediates the proliferation-quiescence decision in the subsequent G1 via p21 expression. Nat. Commun. 8, 14728 (2017).

38. Tan, I.-L. et al. Targeting the non-coding genome and temozolomide signature enables CRISPR-mediated glioma oncolysis. Cell Rep. 42, 113339 (2023).

39. Chen, X. & Calvisi, D. F. Hydrodynamic Transfection for Generation of Novel Mouse Models for Liver Cancer Research. Am. J. Pathol. 184, 912–923 (2014).

40. Wang, X. et al. Preparation of selective organ-targeting (SORT) lipid nanoparticles (LNPs) using multiple technical methods for tissue-specific mRNA delivery. Nat. Protoc. 18, 265–291 (2023).

